# HIF-1α is Required to Differentiate the Neonatal Macrophage Secretome from Adults

**DOI:** 10.1101/2024.04.24.591000

**Authors:** Amanda Becker, Mallory Filipp, Connor Lantz, Kristofor Glinton, Edward B. Thorp

**Author notes:** Correspondence &.

## Abstract

The immune response to stress diverges with age, with neonatal macrophages implicated in tissue regeneration versus tissue scarring and maladaptive inflammation in adults. Integral to the macrophage stress response is the recognition of hypoxia and pathogen-associated molecular patterns (PAMPs), which are often coupled. The age-specific, cell-intrinsic nature of this stress response remains vague. To uncover age-defined divergences in macrophage crosstalk potential after exposure to hypoxia and PAMPs, we interrogated the secreted proteomes of neonatal versus adult macrophages via non-biased mass spectrometry. Through this approach, we newly identified age-specific signatures in the secretomes of neonatal versus adult macrophages in response to hypoxia and the prototypical PAMP, lipopolysaccharide (LPS). Neonatal macrophages polarized to an anti-inflammatory, regenerative phenotype protective against apoptosis and oxidative stress, dependent on *hypoxia inducible transcription factor-1α* (*HIF-1α).* In contrast, adult macrophages adopted a pro-inflammatory, glycolytic phenotypic signature consistent with pathogen killing. Taken together, these data uncover fundamental age and *HIF-1α* dependent macrophage programs that may be targeted to calibrate the innate immune response during stress and inflammation.

## 1. Introduction

Macrophages are innate immune cells with vast functional diversity, plasticity, and presence in most human tissues [1]. They regulate normal physiologic development and tissue homeostasis and are integral mediators in numerous pathophysiologic states to promote tissue repair, inflammation and its resolution, pathogen clearance, immunity, and tissue regeneration [2,3,4]. Previous studies have identified ontological differences in macrophage development contributing to distinct macrophage populations as well as unique gene expression profiles in tissue resident and non-resident macrophages [5,6]. These studies have led to a greater understanding of the developmental origins, differentiation, maintenance, replacement, and functional differences between unique macrophage populations [7,8,9]. However, vast knowledge deficits remain regarding age-specific macrophage phenotypes in response to common physiologic stressors.

Innate immune cell activation by PAMPs such as endotoxin, as well as cellular and tissue hypoxia, are hallmarks of mammalian sepsis regardless of age [10,11,12,13]. Despite sepsis accounting for approximately 20% of annual global deaths with more than 30% of these deaths occurring in children less than 5 years of age, age-specific macrophage responses to hypoxia and endotoxin stimulation remain vague [14,15]. Macrophages in response to hypoxia and LPS activate hypoxia inducible transcription factors (HIFs), a family of transcription factors with isoform-specific functions varying in disparate cell types, to respond to the altered metabolic demands of low oxygen and nutrient supply [16,17,18]. In adult macrophages, the HIF-1α isoform promotes the transcription of genes required for heightened glycolytic metabolism, induction of macrophage mobilization to hypoxic tissue microenvironments, phagocytosis, and upregulation of inflammatory cytokine production [19,20]. In contrast, the downstream effects of neonatal macrophage HIF-1α stabilization in the setting of hypoxia and/or LPS stimulation remain unknown. Uncovering age-specific differences in macrophage and HIF-1α mediated responses to these stimuli may reveal new therapeutic targets to reduce sepsis-related mortality.

To investigate differences in phenotypic plasticity between neonatal and adult macrophages under hypoxic stress and/or LPS stimulation, we interrogated the secretomes of neonatal and adult bone marrow-derived macrophages (BMDMs) from wildtype mice exposed to normoxia (FiO2 0.21), hypoxia (FiO2 0.01), normoxia (FiO2 0.21) + LPS, and hypoxia (FiO2 0.01) + LPS. We then utilized the aforementioned conditions to study differences in the secretomes of neonatal macrophages genetically depleted of *HIF-1α* compared to wildtype neonatal macrophages to uncover neonatal HIF-1α mediated responses to hypoxic stress and/or LPS stimulation. Literature review was performed to delineate known biological functions for the most highly secreted proteins from each BMDM population and experimental condition. Weighted correlation protein network analyses (WCNA) were performed to cluster significantly expressed proteins unique to each macrophage population and experimental condition. Gene ontology was performed to identify upregulated gene pathways within each protein cluster. Herein we have uncovered distinct, age-specific differences in the secretomes and phenotypes of neonatal and adult BMDMs in response to common physiologic stressors such as hypoxia and LPS.

## 2. Materials & Methods

### Neonatal and Adult Mouse Bone Marrow-Derived Macrophage (BMDM) Isolation

C57BL/6 (wildtype) and *LysMcre^+/+^HIF-1α^fl/fl^* mice were euthanized by cervical dislocation after CO_2_ asphyxiation in accordance with IACUC and AVMA guidelines on euthanasia. Both male and female sexes, in equal proportion, were utilized for BMDM isolation. For adult BMDM isolation, bone marrow cells were harvested from the tibias and femurs of mice by cutting both metaphyses and flushing cells from the bone marrow with 5 mL DMEM. For neonatal BMDM isolation, tibias and femurs were placed in a sterile mortar containing 5 mL DMEM and crushed with a pestle to release cells from the bone marrow. The released cells were then passed through a 70 µm filter to remove bone fragments and collagen debris. For both adult and neonatal BMDM isolation, red blood cells were removed by lysis (BioLegend) and filtered through a 70 µm filter. Bone marrow cells were cultured in tissue culture-treated 10 cm Petri dishes (Fisherbrand) for 7 days with DMEM containing 20% L929-cell conditioned media, 10% fetal bovine serum, 1% penicillin-streptomycin, 1% sodium pyruvate, and 1% L-glutamine. A half media change was performed on day 3 of culture. On culture day 7, media was removed, adherent BMDMs were washed with PBS, and 1 mL of cold Cellstripper (Corning) was added to the dish. After 15 minutes of incubation at room temperature, BMDMs were removed and collected by gentle pipetting, washed with DMEM containing 10% FBS, 1% penicillin-streptomycin, 1% sodium pyruvate, and 1% L-glutamine, counted, and seeded to tissue culture-treated plates for experiments.

### Hypoxia and LPS BMDM Treatment

BMDMs were seeded in 24-well plates (Falcon) at a density of 0.24 x 10^6^ cells/well in 1 mL DMEM containing 10% FBS, 1% penicillin-streptomycin, 1% sodium pyruvate, and 1% L-glutamine and allowed to adhere overnight. Prior to hypoxia and/or LPS treatment, the aforementioned media was removed and BMDMs were washed three times with PBS after which 1 mL of serum-free DMEM without phenol red containing 1% penicillin-streptomycin, 1% sodium pyruvate, and 1% L-glutamine was added to each well. For experimental conditions in which LPS stimulation was performed, LPS was added to each well at a concentration of 100 ng/mL. BMDMs exposed to extreme hypoxia (1% O_2_, 94% N_2_, and 5% CO_2_) were placed in a temperature- and humidity-controlled hypoxic chamber (O_2_ Control In Vitro Glove Box, Coy Laboratory Products) for 6 hours. BMDM supernatant was then collected within the hypoxic chamber to minimize reoxygenation effects. BMDMs were labeled with Zombie Aqua Fixable Viability Dye in the hypoxic chamber, fixed, and analyzed by flow cytometry to measure cell death. BMDMs cultured in a temperature- and humidity-controlled tissue culture incubator supplied with room air (21% O_2_, 5% CO_2_) were used as normoxic controls. Three replicates were used for each experimental condition. Total time for hypoxic, normoxic, and/or LPS exposure was 6 hours for each replicate.

### BMDM Secretome Harvest

BMDM supernatant was filtered using a 0.2 µm filter. The filtered supernatant was then concentrated using a centrifugal filter unit with a 3 kDA cutoff at 4°C (Amicon Ultra-4 Centrifugal filters, Millipore, USA). Concentrated supernatant was stored in 200 µL aliquots at −80°C until further analysis.

### Flow Cytometry

BMDM cell counts and viability were evaluated using flow cytometry. BMDMs were stained with Zombie Aqua Fixable Viability Dye (1:1000; BioLegend) in PBS for 15 minutes at room temperature in the dark. Stained BMDMS were spun down and washed with FACS buffer and then fixed with 1:1 FACS buffer and 2% paraformaldehyde. Flow cytometric analyses were performed on a LSRFortessa X-20 Cell Analyzer (BD Biosciences). Data were analyzed using FlowJo software.

### Mass Spectrometry

1. **(A) In-Solution Digestion (ISD)**-ISD was performed using cold Acetone/TCA precipitation as per Northwestern University (NU) Mass Spectrometry Core’s protocol. Pellets were resuspended in 100 µL of 8M Urea/0.4 M AmBic and 4 µL of 100 mM dithiothreitol (DTT) prepared in H_2_O was then added and incubated at 50 °C for 45 min. Next, 6 µL of Indole-3-Acetic Acid (IAA) in H_2_O was added followed by incubation at room temperate in the dark for 45 min, and then 290 µL of AmBic was added. MS-grade trypsin (Promega, Madison WI) was added at an enzyme-to-substrate ratio of 1:50 for overnight digestion at 37 ⁰C. Prior to MS analysis, samples were dried and de-salted using C18 spin columns as per NU Mass Spectrometry Core’s protocol.
2. **(B) LC-MS/MS Analysis**-Peptides were analyzed by LC-MS/MS using a Dionex UltiMate 3000 Rapid Separation nanoLC coupled to a Q-Exactive HF (QE) Quadrupole-Obitrap mass spectrometer (Thermo Fisher Scientific Inc, San Jose, CA). Samples were loaded onto the trap column of 150 μm internal diameter (ID) x 2 cm length in-house packed with 3 μm ReproSil-Pur® beads and the analytical column of 75 μm internal diameter (ID) x 10.5 cm length PicoChip column packed with 3 μm particle size (New Objective, Inc. Woburn, MA). All fractions were eluted from the analytical column at a flow rate of 300 nL/min using an initial gradient of 5% B from 0 to 5 min, ramped to 40% over 100 min, then ramping up to 90% B for 7 min, holding 90% B for 3 min, followed by re-equilibration of 5% B at 10 min with a total run time of 120 min. Mass spectra were acquired by Xcalibur software operating in data-dependent acquisition mode, switching between full scan MS1 (60,000 resolution, 50 ms maximum injection time) and MS2 (30,000 resolution, 100 ms maximum injection time). MS2 spectra were obtained by high-energy collision dissociation (HCD) fragmentation using 30% normalized collision energy.
3. **(C) Data Analysis**-Scaffold 5 software (https://www.proteomesoftware.com/products/scaffold-5) was used to analyze proteomics data unless explicitly stated. Contaminants (10 of 202 identified proteins) were excluded from all statistical analyses. The Normalized Total Spectra for each identified protein was averaged across experimental replicates for statistical analyses using unpaired *t* test and multivariate ANOVA tests with a significance level of <0.05 and false discovery rate 1%. Evaluation of Gene Ontology (GO) biological processes and protein set enrichment analyses (PSEA-quant) were performed using Scaffold 5. Protein network and pathway analyses were performed as below.

### R Analyses

Using an inductive approach, a WCNA was used to cluster all 192 annotated proteins in R v4.1.1 using the WGCNA package (v1.72-1). Modules, or clusters, of proteins were defined based on their weighted correlations to each other. Soft power for protein hierarchical clustering was chosen at the minimum value which resulted in a scale free topology model R^2^>0.85 with mean connectivity minimized. Modules were visualized using igraph (v1.5.1) and for simplicity, edges were only shown if the absolute value correlation between proteins was greater than 0.001. Main hub proteins were identified using chooseTopHubInEachModule() within the WGCNA package. Pathway analyses on the modules were also performed using g:Profiler (version: e109_eg56_p17_1d3191d) [21].

### General Quantification and Statistical Analyses

Statistical analyses were performed with GraphPad Prism 9 software (Graphpad Software). Comparisons between groups were performed using a two-tailed, unpaired *t* test with a significance level of <0.05 and 95% confidence interval. For comparisons of more than two variables, a one-way or two-way ANOVA was used with a 95% confidence interval. When necessary, a Tukey test was used to correct for multiple comparisons. *P* values < 0.05 were considered statistically significant (*, *P* < 0.05; **, *P* < 0.01; ***, *P* < 0.001). Analyses labeled “ns” were not statistically significant.

## 3. Results

### Hypoxia and LPS stimulate neonatal and adult BMDMs to secrete unique proteomes

A total of 192 distinct proteins were identified via mass spectrometry from the secretomes of BMDMs harvested from neonatal C57BL/6, neonatal *LysMcre^+/+^HIF-1α^fl/fl^*, and adult C57BL/6 mice exposed to either normoxia (FiO2 0.21), hypoxia (FiO2 0.01), normoxia (FiO2 0.21) + LPS, or hypoxia (FiO2 0.01) + LPS. Each experimental condition stimulated neonatal and adult BMDM populations to secrete unique proteomes (**Figures 1 and 2**). Hypoxia and LPS, individually, as well as hypoxia + LPS stimulation induced neonatal and adult BMDM populations to secrete distinct proteins compared to normoxic controls (**Figures 1 and 2**). Under normoxic conditions, the proteomes of neonatal C57BL/6 and neonatal *LysMcre^+/+^HIF-1α^fl/fl^* as well as adult C57BL/6 and neonatal *LysMcre^+/+^HIF-1α^fl/fl^* shared more overlapping proteins than the proteomes of neonatal and adult C57BL/6 (**Figure 1A**). Hypoxia induced more robust protein secretion from neonatal C57BL/6 BMDMs compared to neonatal *LysMcre^+/+^HIF-1α^fl/fl^* and adult C57BL/6 BMDMs (**Figure 1B**). LPS, under both normoxic and hypoxic conditions, stimulated greater protein secretion from adult C57BL/6 BMDMs compared to both neonatal BMDM populations (**Figure 2A and 2B**). Live/dead staining, performed via flow cytometry, revealed >97% survival for neonatal and adult BMDMs exposed to hypoxia, LPS, or hypoxia + LPS, suggesting cell death in response to LPS and/or hypoxia was unlikely to be contributing to the differing secreted proteomes (**Supplemental Figure 1**). WCNA identified unique and overlapping clusters of significantly expressed proteins for each macrophage population and experimental condition (**Figure 3A and 3C, Figure 4A and 4C, Supplemental Figure 2**). Ontology analysis identified the most common upregulated gene pathways within each cluster (**Figures 3B and 4C**). Volcano plots were constructed to detect significant differences in BMDM protein secretion between neonatal C57BL/6, neonatal *LysMcre^+/+^HIF-1α^fl/fl^*, and adult C57BL/6 mice exposed to normoxia (**Figure 5**), hypoxia (**Figure 6**), normoxia + LPS (**Figure 7**), or hypoxia + LPS (**Figure 8**). BMDM population specific differences in protein secretion under normoxic, hypoxic, and LPS conditions (**Supplemental Figures 3, 4, and 5**) were also interrogated. Known biological functions for proteins secreted from each BMDM population and experimental condition are presented in **Supplemental Table 1**.

**Figure 1:**
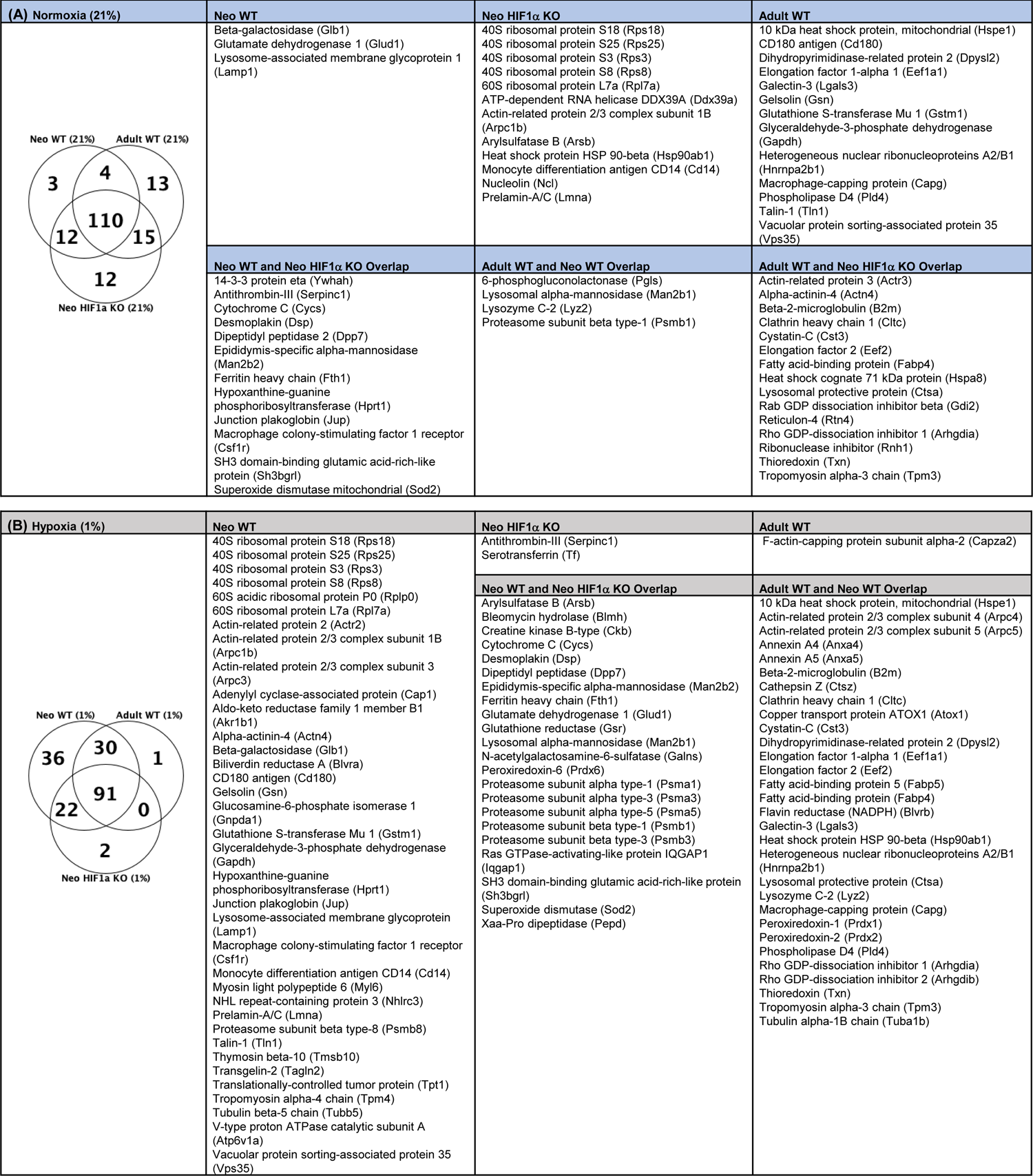
Neonatal and adult macrophages secrete unique proteomes under normoxic and hypoxic conditions. (A) Unique and overlapping proteins secreted from neonatal wildtype, neonatal *LysMcre^+/+^HIF-1α^fl/fl^* (HIF-1α KO), and adult wildtype BMDMs under normoxic conditions (FiO2 0.21). (B) Unique and overlapping proteins secreted from neonatal wildtype, neonatal *LysMcre^+/+^HIF-1α^fl/fl^* (HIF-1α KO), and adult wildtype BMDMs with hypoxic stimulation (FiO2 0.01).

**Figure 2:**
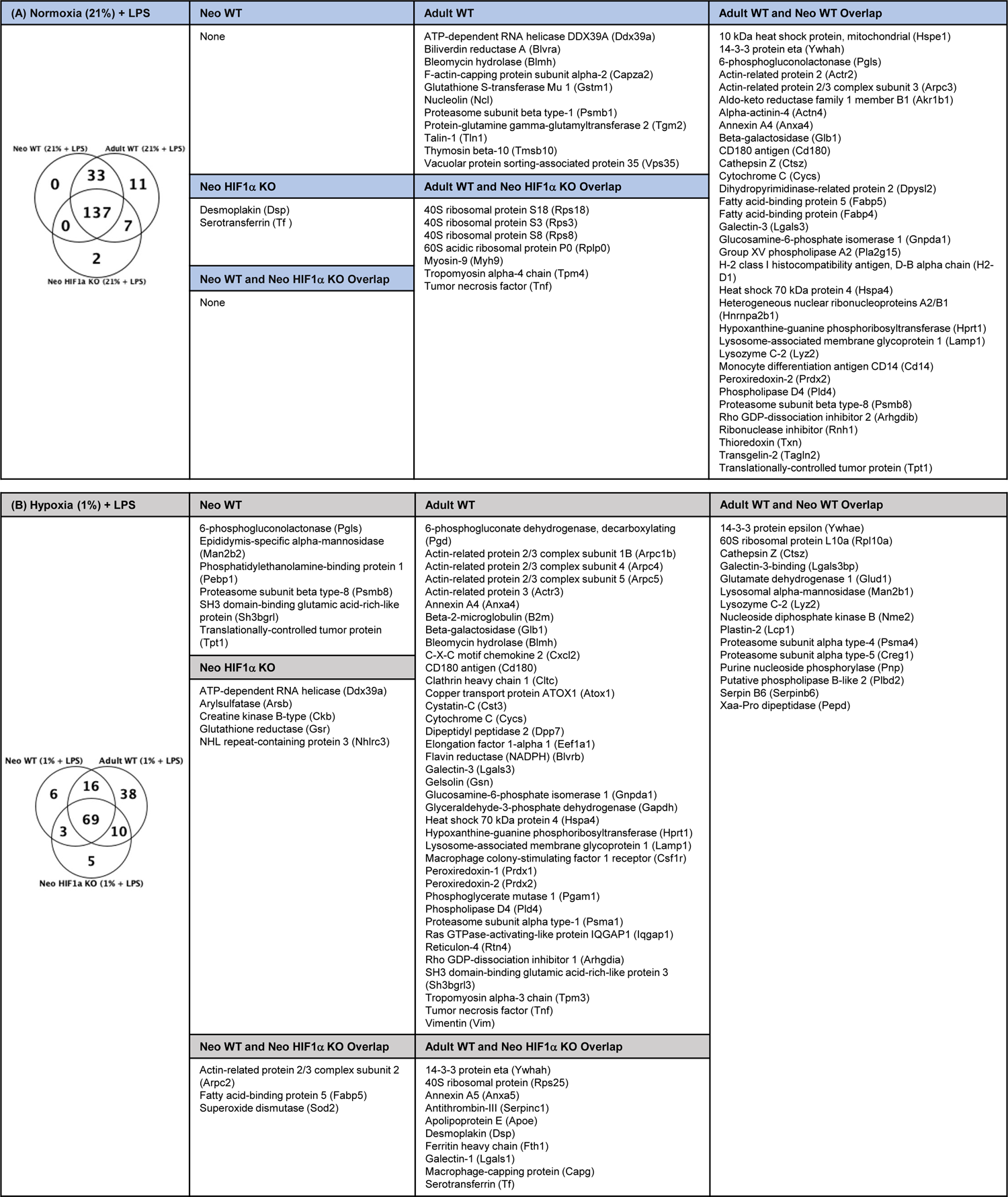
Neonatal and adult macrophages secrete unique proteomes with LPS stimulation alone and with LPS stimulation in the setting of hypoxia. (A) Unique and overlapping proteins secreted from neonatal wildtype, neonatal *LysMcre^+/+^HIF-1α^fl/fl^* (HIF-1α KO), and adult wildtype BMDMs under LPS stimulation in the setting of normoxia (FiO2 0.21) (B) Unique and overlapping proteins secreted from neonatal wildtype, neonatal *LysMcre^+/+^HIF-1α^fl/fl^* (HIF-1α KO), and adult wildtype BMDMs with LPS stimulation in the setting of hypoxia (FiO2 0.01).

**Figure 3:**
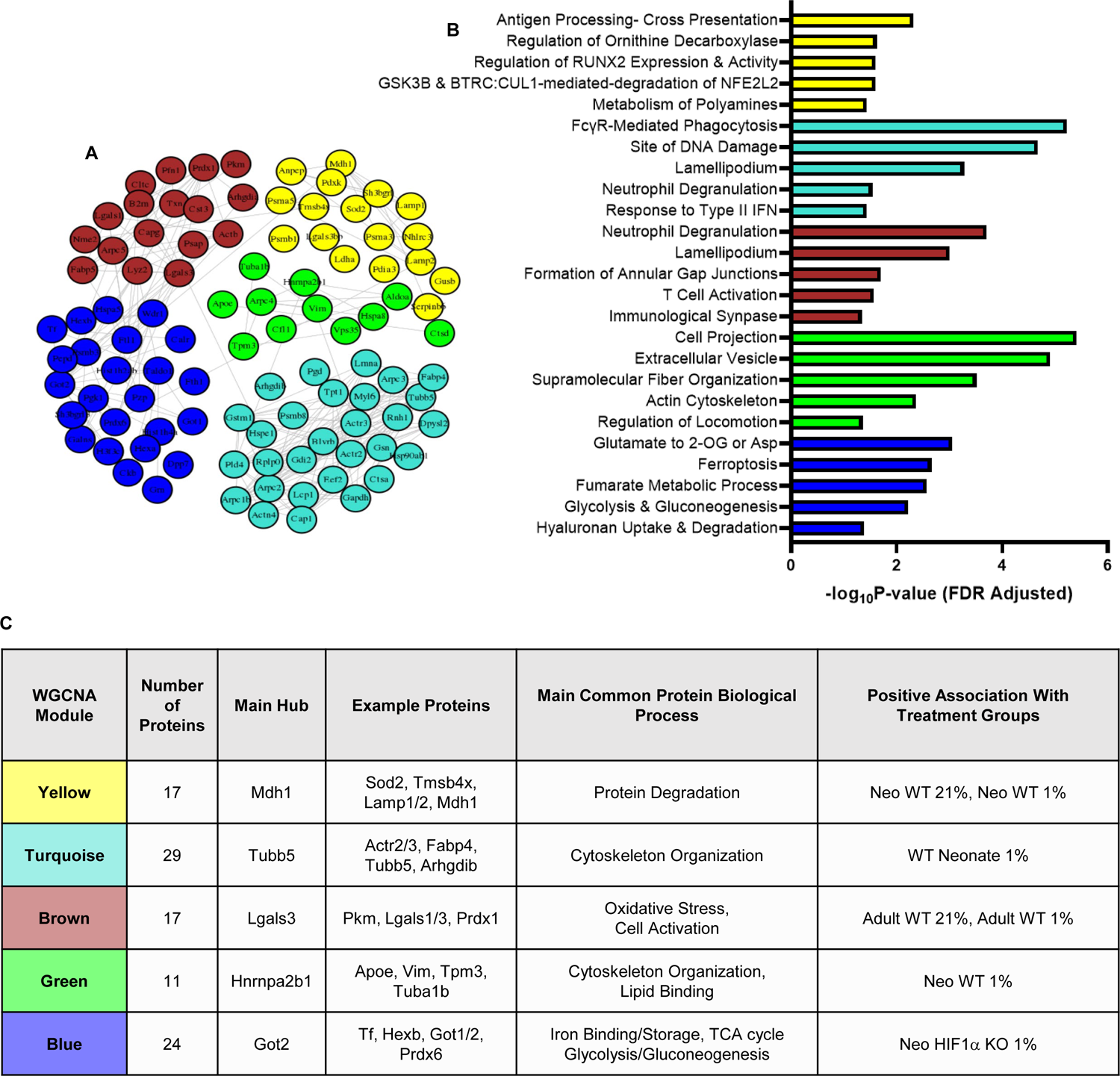
Neonatal and adult macrophages secrete unique protein clusters with specific upregulated gene pathways under normoxic and hypoxic conditions. (A) Adjacency network map of secreted proteins identified by proteomics analysis, color coded by cluster assignment hierarchical clustering-based nearness or co-expression of proteins. For clarity, only proteins associated with a cluster (98/192) and edges representing correlations greater than 0.001 are displayed. (B) Overrepresented, nonredundant upregulated gene pathways in each cluster with false discovery rate corrected *P* values. (C) Weighted gene co-expression network analysis summary of extracted protein clusters including protein cluster size, main hub protein, most common protein biological process, and protein cluster association with macrophage population and experimental condition.

**Figure 4:**
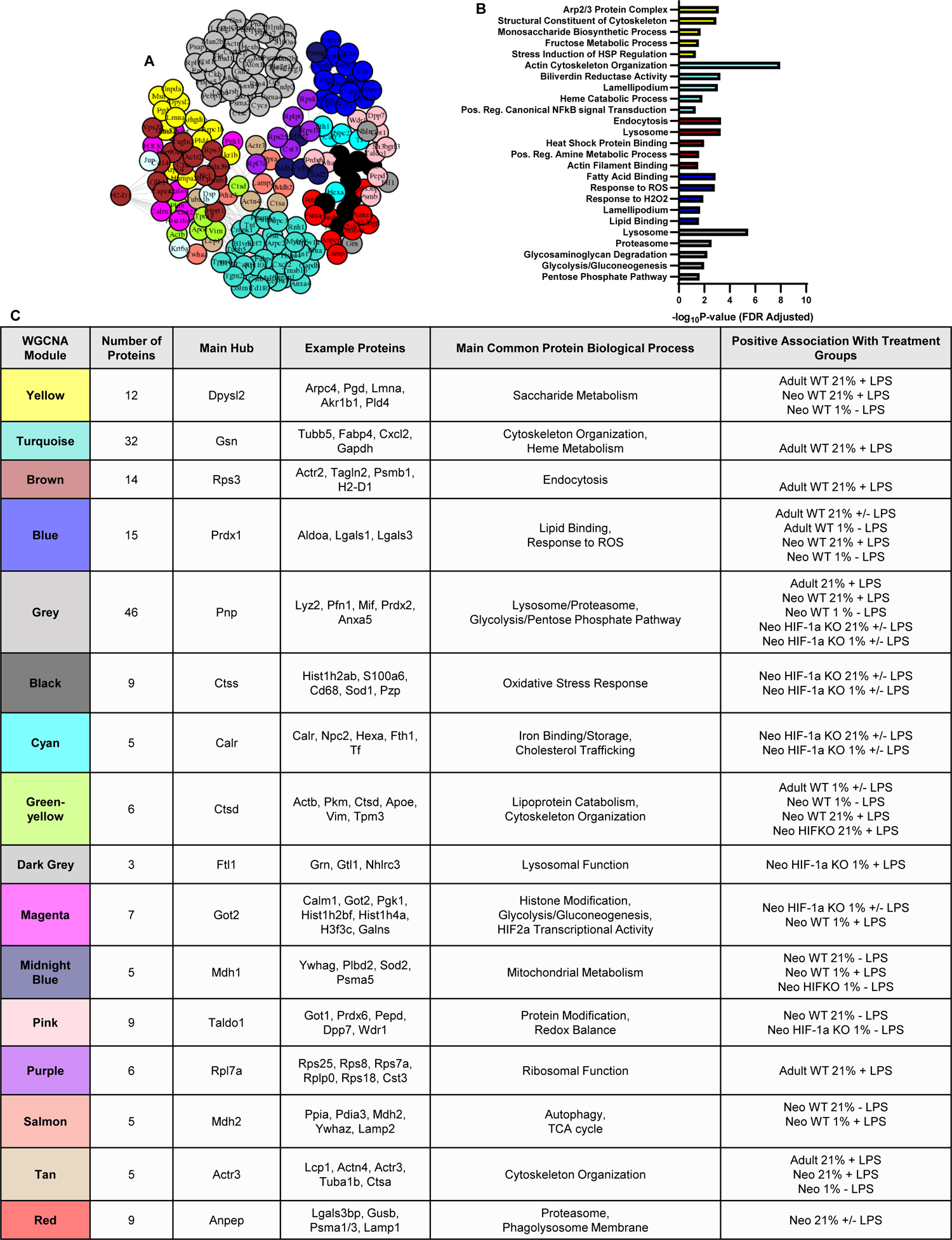
Neonatal and adult macrophages secrete unique protein clusters with specific upregulated gene pathways with LPS stimulation alone and with LPS stimulation plus hypoxia. (A) Adjacency network map of secreted proteins identified by proteomics analysis, color coded by cluster assignment hierarchical clustering-based nearness or co-expression of proteins. For clarity, only proteins associated with a cluster (98/192) and edges representing correlations greater than 0.001 are displayed. (B) Overrepresented, nonredundant upregulated gene pathways in each cluster with false discovery rate corrected *P* values. (C) Weighted gene co-expression network analysis summary of extracted protein clusters including protein cluster size, main hub protein, most common protein biological process, and protein cluster association with macrophage population and experimental condition.

**Figure 5:**
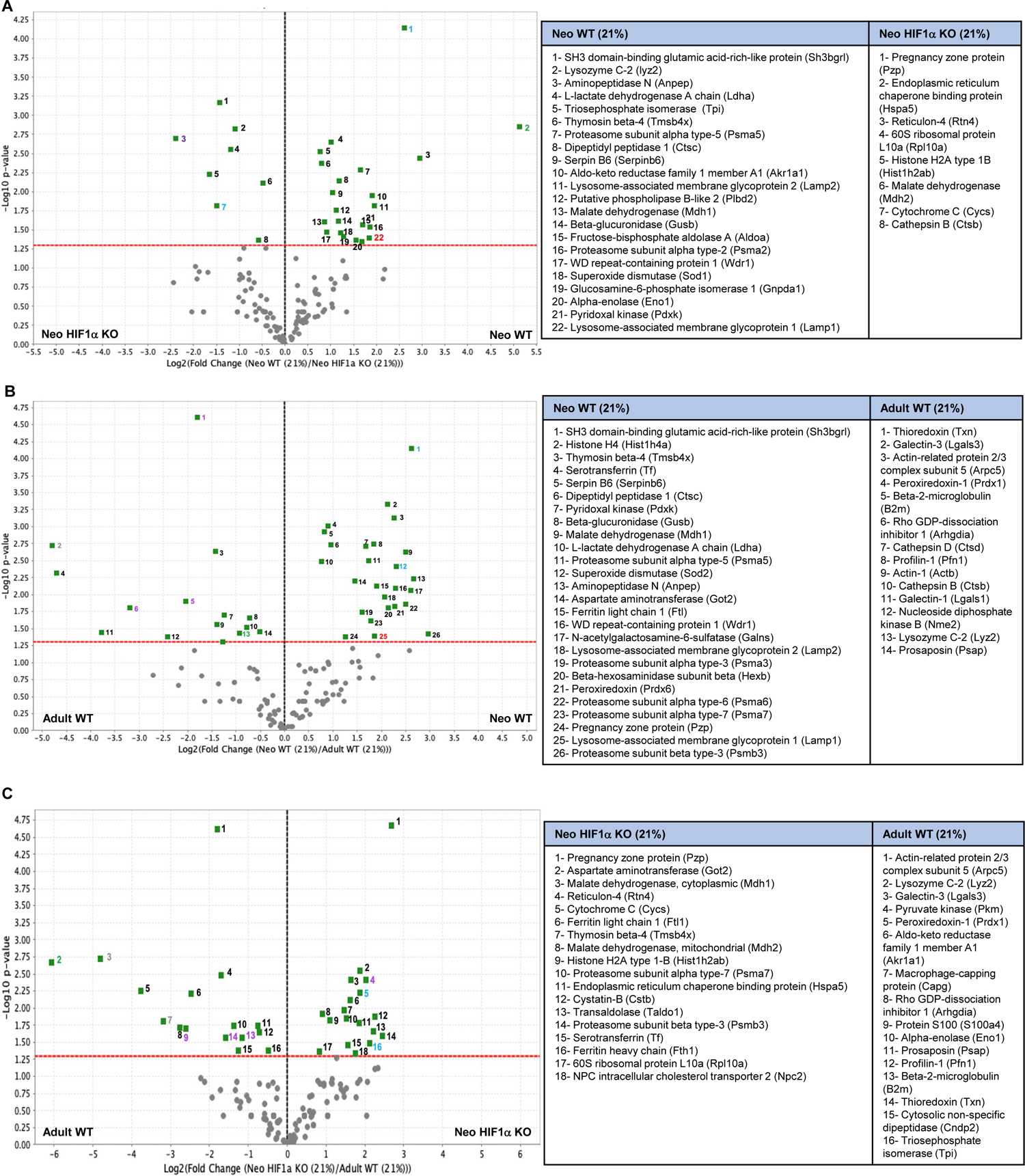
Neonatal wildtype (Neo WT), neonatal *LysMcre^+/+^HIF-1α^fl/fl^* (Neo HIF-1α KO), and adult wildtype (Adult WT) macrophages secrete significantly different proteins in the setting of normoxia. Volcano plots depicting significant differences in protein secretion between (A) Neo WT and Neo HIF-1α KO BMDMS, (B) Neo WT and Adult WT BMDMs, and (C) Neo HIF-1α KO and Adult BMDMs. For all volcano plots the red dashed line represents the significance threshold and the black dashed line represents zero-fold change between groups utilizing t-tests with a significance level of < 0.05. Protein colored numbering: black= proteins secreted from all macrophage populations, purple= proteins secreted from neo HIF-1α KO and adult WT BMDMs, blue= proteins secreted from neo WT and neo HIF-1α KO BMDMs, green= proteins secreted from neo WT and adult WT BMDMs, red= proteins secreted only from neo WT BMDMs, yellow=proteins secreted only from neo HIF-1α KO BMDMs, grey= proteins secreted only from adult WT BMDMs.

**Figure 6:**
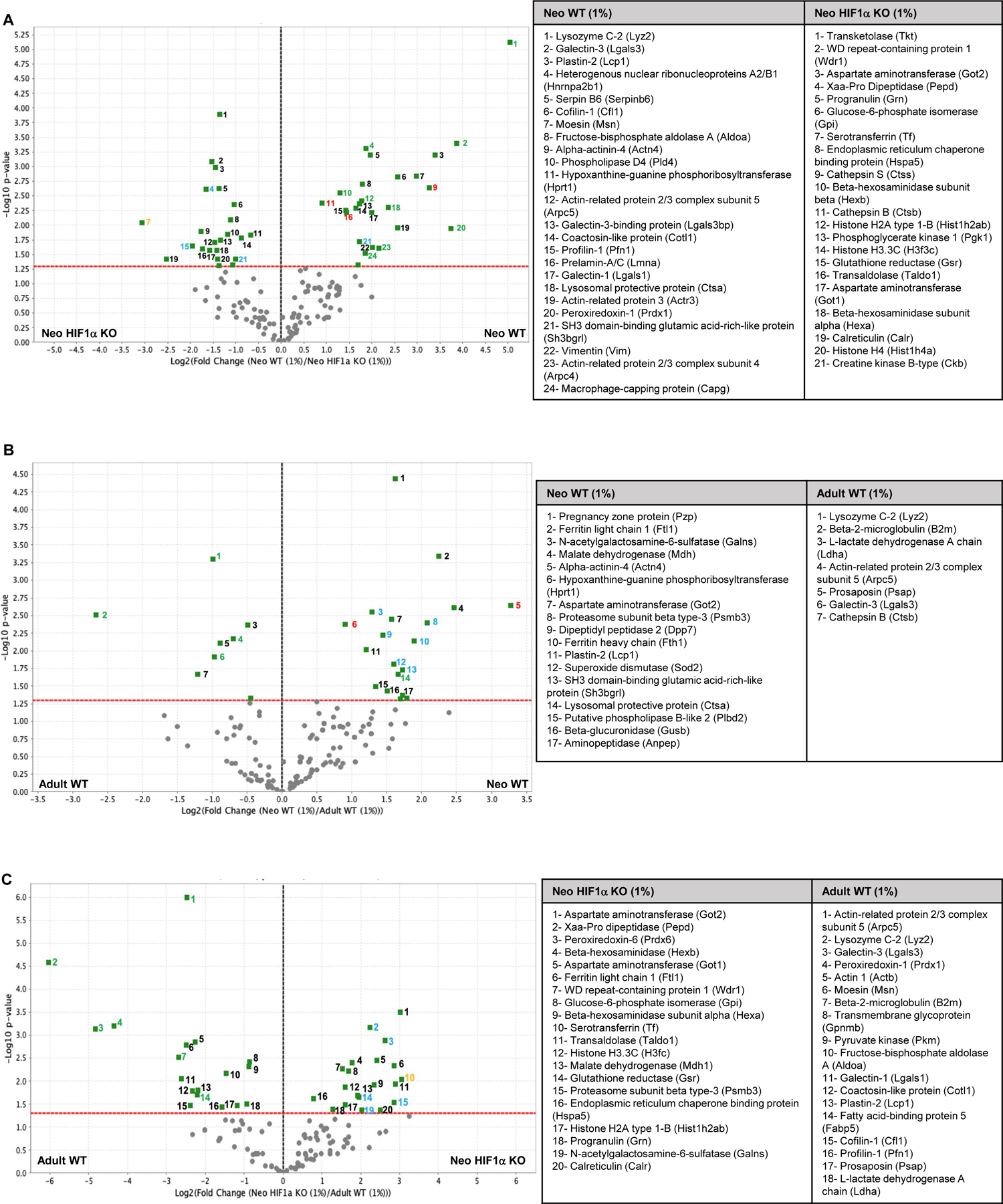
Neonatal wildtype (Neo WT), neonatal *LysMcre^+/+^HIF-1α^fl/fl^* (Neo HIF-1α KO), and adult wildtype (Adult WT) macrophages secrete significantly different proteins with hypoxic stimulation. Volcano plots depicting significant differences in protein secretion between (A) Neo WT and Neo HIF-1α KO BMDMS, (B) Neo WT and Adult WT BMDMs, and (C) Neo HIF-1α KO and Adult BMDMs. For all volcano plots the red dashed line represents the significance threshold and the black dashed line represents zero-fold change between groups utilizing t-tests with a significance level of < 0.05. Protein colored numbering: black= proteins secreted from all macrophage populations, purple= proteins secreted from neo HIF-1α KO and adult WT BMDMs, blue= proteins secreted from neo WT and neo HIF-1α KO BMDMs, green= proteins secreted from neo WT and adult WT BMDMs, red= proteins secreted only from neo WT BMDMs, yellow= proteins secreted only from neo HIF-1α KO BMDMs, grey= proteins secreted only from adult WT BMDMs.

**Figure 7:**
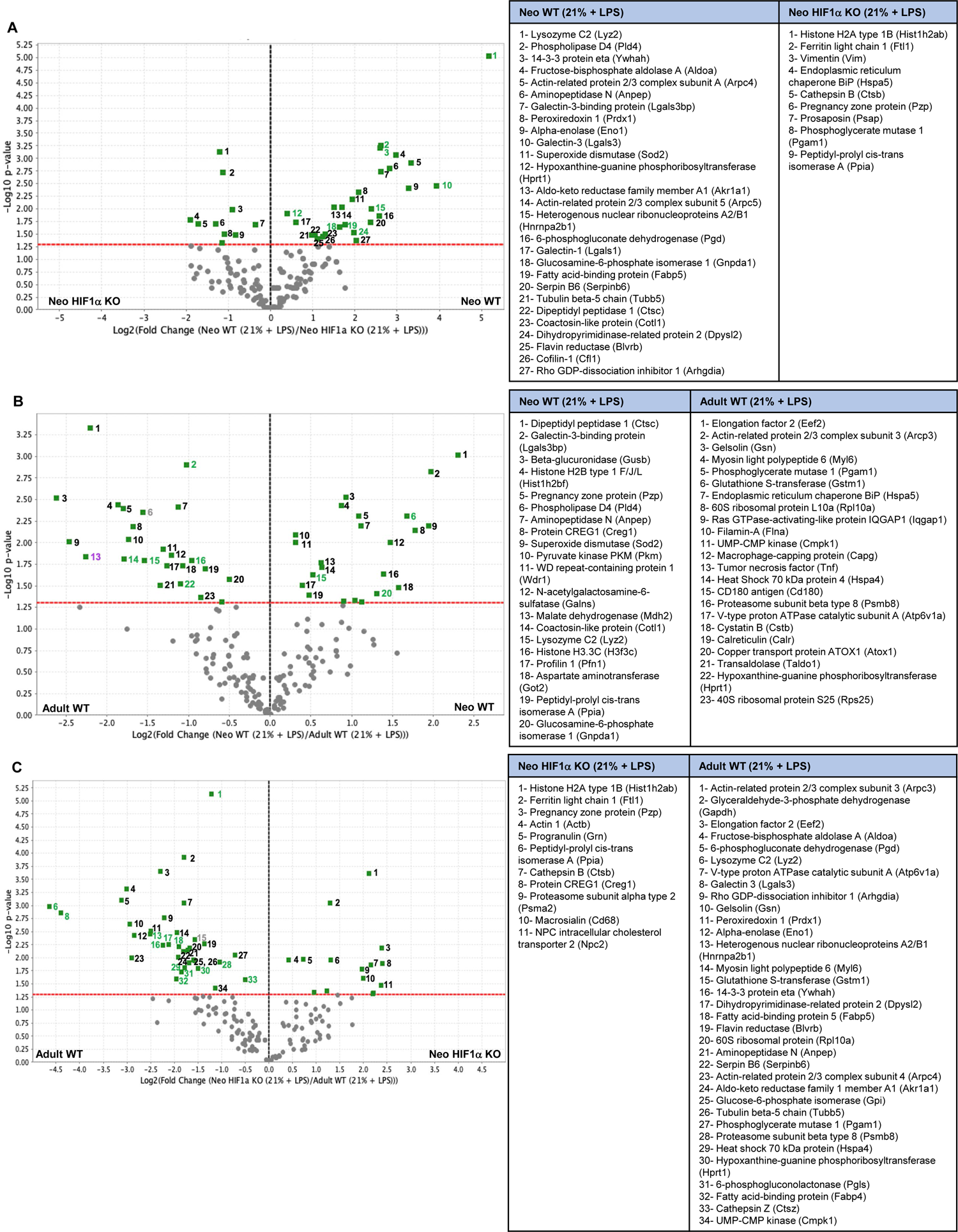
Neonatal wildtype (Neo WT), neonatal *LysMcre^+/+^HIF-1α^fl/fl^* (Neo HIF-1α KO), and adult wildtype (Adult WT) macrophages secrete significantly different proteins with LPS stimulation alone. Volcano plots depicting significant differences in protein secretion between (A) Neo WT and Neo HIF-1α KO BMDMS, (B) Neo WT and Adult WT BMDMs, and (C) Neo HIF-1α KO and Adult BMDMs. For all volcano plots the red dashed line represents the significance threshold and the black dashed line represents zero-fold change between groups utilizing t-tests with a significance level of < 0.05. Protein colored numbering: black= proteins secreted from all macrophage populations, purple= proteins secreted from neo HIF-1α KO and adult WT BMDMs, blue= proteins secreted from neo WT and neo HIF-1α KO BMDMs, green= proteins secreted from neo WT and adult WT BMDMs, red= proteins secreted only from neo WT BMDMs, yellow= proteins secreted only from neo HIF-1α KO BMDMs, grey= proteins secreted only from adult WT BMDMs.

**Figure 8:**
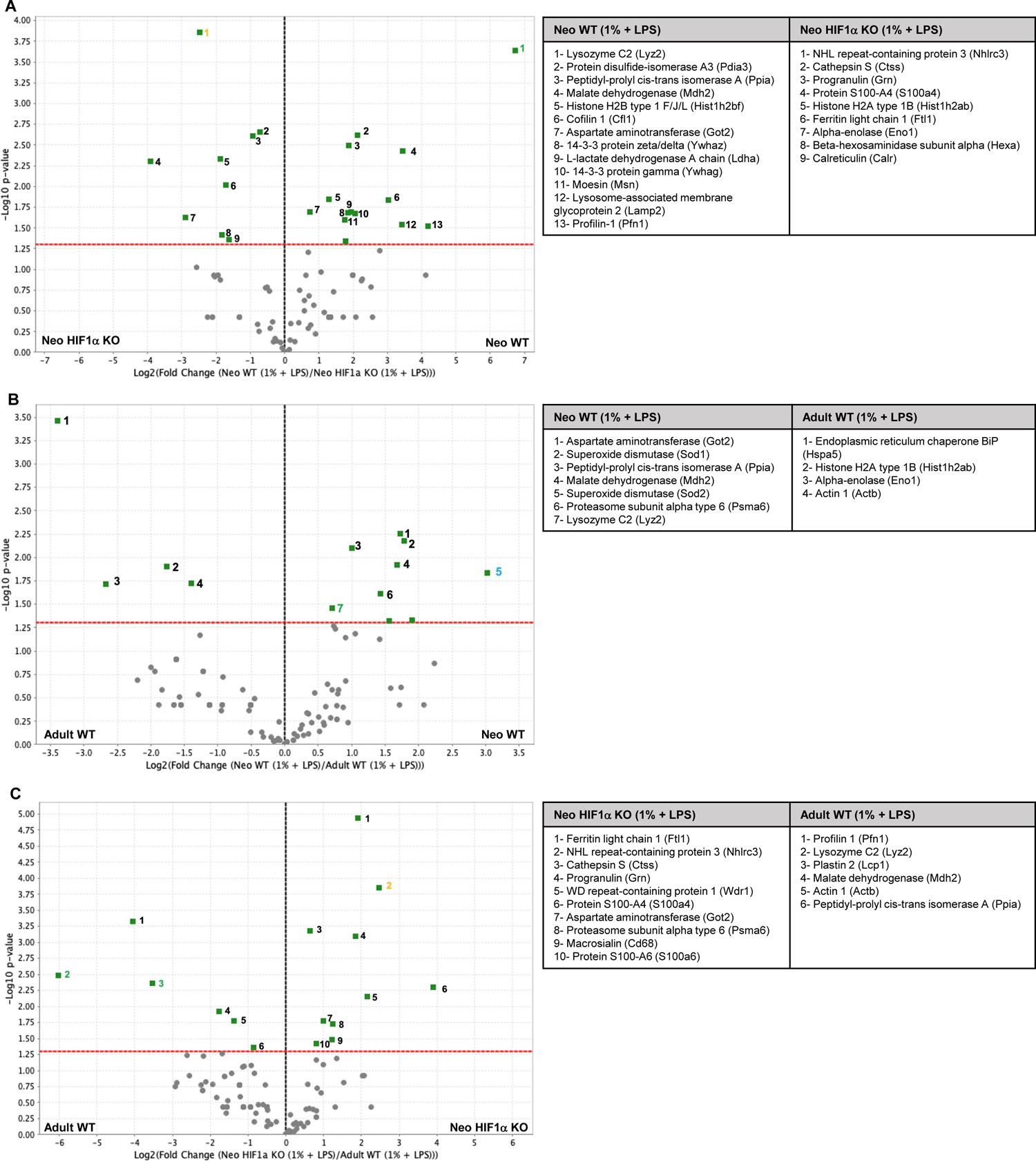
Neonatal wildtype (Neo WT), neonatal *LysMcre^+/+^HIF-1α^fl/fl^* (Neo HIF-1α KO), and adult wildtype (Adult WT) macrophages secrete significantly different proteins with LPS stimulation plus hypoxia. Volcano plots depicting significant differences in protein secretion between (A) Neo WT and Neo HIF-1α KO BMDMS, (B) Neo WT and Adult WT BMDMs, and (C) Neo HIF-1α KO and Adult BMDMs. For all volcano plots the red dashed line represents the significance threshold and the black dashed line represents zero-fold change between groups utilizing t-tests with a significance level of < 0.05. Protein colored numbering: black= proteins secreted from all macrophage populations, purple= proteins secreted from neo HIF-1α KO and adult WT BMDMs, blue= proteins secreted from neo WT and neo HIF-1α KO BMDMs, green= proteins secreted from neo WT and adult WT BMDMs, red= proteins secreted only from neo WT BMDMs, yellow= proteins secreted only from neo HIF-1α KO BMDMs, grey= proteins secreted only from adult WT BMDMs.

### Hypoxia induces neonatal wildtype BMDMs to secrete proteins involved in anti-inflammatory immune modulation, regeneration, and protection from apoptosis

Under normoxic conditions, biological activities of the most highly secreted proteins from neonatal C57BL/6 BMDMs included protection against oxidative stress, regulation of chromosome structure and function, actin polymerization, cell motility, protein catabolism, iron transport and metabolism, anaerobic metabolism, glycolysis, lysosome stabilization, cell death inhibition, cytoskeleton organization, aldehyde and ketone metabolism, carbohydrate metabolism, citric acid cycle (TCA), gluconeogenesis, and ROS inhibition (**Figure 5A and 5B, Supplemental Table 1**). When exposed to hypoxia, neonatal C57BL/6 BMDMs continued to secrete proteins involved in protein catabolism, iron transport, cell death inhibition, glycolysis, TCA cycle, gluconeogenesis, and protection against oxidative stress (**Figure 6A and 6B, Supplemental Table 1**). Gene ontology also identified upregulated gene pathways involved in protection against oxidative stress and cellular apoptosis in neonatal C57BL/6 BMDMs under both normoxic and hypoxic conditions (**Figure 3B**). Hypoxia altered neonatal C57BL/6 BMDM gene expression and protein production to secrete proteins with biological functions specific to bacterial cell death, anti-inflammatory immune modulation, iron storage and detoxification, cell-to-cell communication, cell growth and differentiation, regeneration, angiogenesis, macrophage activation, actin cross-linking for cell adhesion/motility/antigen receptor signaling, RNA metabolism, cell proliferation, cytokinesis, cytoskeleton organization, purine nucleotide production, actin polymerization, arachidonic acid metabolism, mitosis, DNA replication/transcription, and chromatin organization (**Figure 3B, Figure 6A and 6B, Supplemental Table 1**).

### Hypoxia stimulated neonatal BMDMs genetically depleted of *HIF-1α* to secrete proteins involved in glycolysis and induced an adult-like macrophage response to stress

Neonatal *LysMcre^+/+^HIF-1α^fl/fl^* BMDMs under normoxic conditions secreted proteins involved in anti-inflammatory immune modulation, protein chaperoning, inhibition of neuroendocrine signaling, protein production, DNA packaging into chromatin, TCA cycle, gluconeogenesis, electron transport chain, aspartate to glutamate interconversion, and iron transportation and storage (**Figure 5A and 5C, Supplemental Table 1**). When exposed to hypoxia, neonatal *LysMcre^+/+^HIF-1α^fl/fl^* BMDMs continued to secrete proteins involved in aspartate to glutamate interconversion, gluconeogenesis, iron transportation and storage, TCA cycle, and protein chaperoning (**Figure 6A and 6C, Supplemental Table 1**). Hypoxia altered neonatal *LysMcre^+/+^HIF-1α^fl/fl^* BMDM protein production resulting in the secretion of proteins engaged in the pentose phosphate pathway, actin filament disassembly, protein catabolism, antioxidant activities, lysosome activation, glycolysis, cytokine secretion, iron regulation, MHCII antigen presentation, apoptosis, and gene transcription (**Figure 6A and 6C, Supplemental Table 1**). Upregulated gene pathways involved in glycolysis and ferroptosis were also identified in neonatal *LysMcre^+/+^HIF-1α^fl/fl^* BMDMs exposed to hypoxia (**Figure 3B**).

### Adult wildtype BMDMs in contrast to neonatal BMDMs had a less robust response to hypoxia with protein secretion focused on bacterial cell death, glycolytic metabolism, lysosome function, and apoptosis

Adult C57BL/6 BMDMs, under normoxic conditions secreted proteins involved in ROS inhibition, antioxidant activities, actin polymerization, cell-to-cell communication, protection from apoptosis, cell growth and differentiation, immune cell activation and tissue invasion, inhibition of cell proliferation, glycolysis, and aldehyde and ketone metabolism (**Figure 5B and 5C, Supplemental Table 1**). Upregulated gene pathways identified with hypoxic activation of adult C57BL/6 BMDMs included neutrophil degranulation and immune activation (**Figure 3B**). New proteins secreted under hypoxic activation included proteins involved in bacterial cell death, protein catabolism, lysosome function, and apoptosis (**Figure 6B and 6C, Supplemental Table 1**).

### LPS stimulation of neonatal wildtype BMDMs under both normoxic and hypoxic conditions induced neonatal BMDMs to secrete a proteome similar to that under hypoxic stress alone

The biological functions of the most highly secreted proteins from neonatal C57BL/6 BMDMs exposed to LPS under normoxic conditions included bacterial cell death, protein catabolism, suppression of inflammatory cytokine production and ROS production, hematopoietic progenitor cell differentiation, mitosis, cell survival, glycolysis, carbohydrate metabolism, actin polymerization, DNA organization, regeneration, antioxidant function, inhibition of apoptosis, heat shock proteins, cell-to-cell communication, cell growth and differentiation, cell proliferation, macrophage activation, purine nucleotide production, aldehyde and ketone metabolism, RNA metabolism, pentose phosphate pathway, lysosome function, TCA cycle, gluconeogenesis, arachidonic acid metabolism, and gene transcription (**Figure 7A and 7B, Supplemental Table 1**). Upregulated gene pathways involved in saccharide metabolism, glycolysis, heat shock protein regulation, and response to oxidative stress were also identified (**Figure 4B**) in neonatal C57BL/6 BMDMs exposed to LPS under normoxic conditions. When exposed to LPS under hypoxic conditions, neonatal C57BL/6 BMDMs continued to secrete proteins involved in bacterial cell death, protein catabolism, TCA cycle, gluconeogenesis, DNA organization, cell survival, inhibition of apoptosis, glycolysis, antioxidant function, and ROS inhibition (**Figure 8A and 8B, Supplemental Table 1**). Hypoxia in addition to LPS activation, modified BMDM protein to include proteins involved in proper protein folding, epigenetic regulation of transcription, cell motility, cytokinesis, amino acid metabolism, cell cycle progression, and HIF-1α stabilization (**Figure 8A and 8B, Supplemental Table 1**).

### Neonatal BMDMs genetically depleted of *HIF-1α* exposed to LPS under normoxic and hypoxic conditions secrete proteins involved in glycolysis, iron regulation, anti-inflammatory immune modulation, and lysosome function. BMDM secretion of proteins involved in the acceleration of cellular aging and MHCII antigen presentation are specific to LPS stimulation under normoxic versus hypoxic conditions, respectively

Neonatal *LysMcre^+/+^HIF-1α^fl/fl^* BMDMs under normoxic conditions and LPS stimulation secreted proteins involved in DNA packaging into chromatin, iron transportation and storage, iron detoxification, microtubule and cytoskeleton stabilization, protein chaperoning, apoptosis, anti-inflammatory immune modulation, lysosome function, glycolysis, protein folding, cell survival, IL-10 production, acceleration of cellular senescence, and autophagy (**Figure 7A and 7C, Supplemental Table 1**). BMDMs exposed to hypoxia and LPS resulted in ongoing secretion of proteins involved in iron transport and storage, iron detoxification, cell survival, IL-10 production, lysosome function, cell differentiation, DNA packaging into chromatin, and glycolysis. With LPS stimulation under both hypoxic and normoxic conditions, neonatal *LysMcre^+/+^HIF-1α^fl/fl^* BMDM upregulated gene pathways involved in iron regulation and glycolysis (**Figure 4B**). New proteins secreted under these conditions included proteins with biological functions specific to protein ubiquination, MHCII antigen presentation, cell motility and invasion, heat shock proteins, amino acid metabolism, and proteolysis (**Figure 8A and 8C, Supplemental Table 1**).

### Adult wildtype BMDMs had a robust response to LPS stimulation under normoxic conditions with protein production focused on glycolysis and glycolytic alternative metabolism, inflammation, macrophage activation, cell invasion, pathogen killing, and protein production

Adult C57BL/6 BMDMs exposed to LPS under normoxic conditions secreted proteins involved in protein synthesis, ribosomal translocation, actin polymerization, glycolysis, pentose phosphate pathway, bacterial cell death, protein catabolism, cell-to-cell communication, cell differentiation, macrophage activation, protection from apoptosis, NADPH oxidase activation, cellular detoxification, antioxidant function, ROS inhibition, RNA metabolism, protein chaperoning, ribosome subunit, integrin binding, cell migration/adhesion/invasion, microtubule assembly, fatty acid transport, nucleic acid biosynthesis, oxidative stress, proinflammatory cytokines, insulin resistance, apoptosis, heat shock proteins, protein folding, toll-like receptor 4, proteolysis, copper transport and homeostasis (**Figure 7B and 7C, Supplemental Table 1).** Significantly upregulated gene pathways involved in saccharide metabolism, heme metabolism, proinflammatory cytokine and chemokine production, inflammasome activation, endocytosis, ribosomal function, lysosome and proteasome activation, glycolysis, and the pentose phosphate pathway were also observed in adult C57BL/6 BMDMs exposed to LPS under normoxic conditions (**Figure 4B**). Hypoxia and LPS exposure led to ongoing secretion of proteins specific to protein chaperoning, glycolysis, cell migration and invasion, actin polymerization, bacterial cell death, heat shock proteins, and protein catabolism (**Figure 8B and 8C, Supplemental Table 1**). The biological functions of newly secreted proteins from adult C57BL/6 BMDMs exposed to LPS under hypoxic conditions included DNA packaging into chromatin, vesicle and organelle movement, TCA cycle, and gluconeogenesis (**Figure 8B and 8C, Supplemental Table 1**).

## 4. Discussion

We sought to uncover age-specific differences in macrophage secretomes in response to common physiologic stressors such as hypoxia and the endotoxin, LPS, as well as *HIF-1α* dependent responses to these stimuli in neonatal macrophages. Collectively, we identified adult and neonatal BMDMs secrete distinct, unique proteomes in response to hypoxia or LPS stimulation. Neonatal wildtype BMDMs assume an anti-inflammatory, regenerative signature protective against apoptosis and oxidative stress in response to hypoxia or LPS stimulation. Interestingly, neonatal BMDMs genetically depleted of *HIF-1α* do not assume a regenerative phenotype in response to hypoxic stress or LPS stimulation. Instead, *HIF-1α* deficient neonatal BMDMs adopt an adult-like response to these stimuli with secretion of proteins involved in glycolysis, lysosome activation, and apoptosis but remain anti-inflammatory in nature. Lastly, although hypoxia induced less robust protein secretion compared to LPS stimulation from adult wildtype macrophages, a similar response was observed with protein secretion focused on pathogen killing, glycolytic metabolism, lysosome activation, and apoptosis. LPS also induced a pro-inflammatory phenotype from adult wildtype BMDMs (**Supplemental Figure 6**). To our knowledge, this is the first study examining neonatal *HIF-1α* dependent and age-specific differences in neonatal versus adult BMDM protein secretory responses to hypoxia and LPS stimulation. The pro-inflammatory, glycolytic, and pathogen killing phenotype we observed in adult wildtype BMDMs under hypoxic stress and LPS stimulation corroborates previous research examining adult BMDM responses to the aforementioned stimuli [10,20,22,23]. As vast knowledge deficits remain regarding neonatal BMDM and neonatal *HIF-1α* dependent responses to common pathophysiologic stimuli, the remainder of our discussion will focus on the unique secretomes of *HIF-1α* expressing and *HIF-1α* deficient neonatal BMDMs.

With all experimental conditions, we observed robust protein secretion involved in glycolytic metabolism, protection against oxidative stress, and inhibition of apoptosis from neonatal wildtype BMDMs. Mammals develop in a hypoxic environment in utero with hypoxia being a known molecular trigger for normal fetal development and glycolysis being the predominant energy source. Increased secretion of glycolytic proteins in response to stress may stem from retained fetal pathways that allow for normal development in the setting of hypoxia [24,25,26,27]. Despite glycolytic metabolism, an anti-inflammatory phenotype was observed in both *HIF-1α* expressing and *HIF-1α* deficient neonatal BMDMs with hypoxic stress or LPS stimulation, suggesting that glycolysis may not induce a pro-inflammatory phenotype in neonatal BMDMs as has previously been described in adult BMDMs [28,29]. The upregulation of genes involved in glycolysis as well as glycolytic protein secretion from *HIF-1α* deficient neonatal BMDMs also suggests that *HIF-1α* may not be a major stimulator of glycolysis in neonatal BMDMs, in contrast to adult BMDMs, or that alternative mechanisms for glycolytic activation exist in neonatal BMDMs [30,31]. Furthermore, lactate, as a byproduct of glycolysis, is a known epigenetic regulator via histone lysine lactylation (Kla) [32,33]. In *HIF-1α* expressing neonatal BMDMs, we observed increased secretion of proteins involved in chromatin organization, chromosome function and structure, gene transcription, and the epigenetic regulation of gene transcription, suggesting a potential role for glycolysis mediated epigenetic regulation of neonatal BMDMs and a possible mechanism for *HIF-1α* mediated neonatal regeneration.

In *HIF-1α* deficient neonatal BMDMs, we observed the upregulation of genes involved in ferroptosis with hypoxic stimulation as well as increased secretion of proteins involved in iron transportation, storage, and detoxification with all experimental conditions suggesting a *HIF-1α* mediated mechanism for protection against iron-dependent cell death. With both hypoxic and LPS stimulation, *HIF-1α* deficient neonatal BMDMs secreted proteins associated with *HIF-2α* transcriptional activity suggesting greater dependence on *HIF-2α* in the absence of *HIF-1α* and possible *HIF-2α* mediated adult-like programming of neonatal BMDMs. Finally, in *HIF-1α* deficient neonatal BMDMs exposed to LPS we observed increased secretion of proteins involved in the acceleration of cellular aging, suggesting *HIF-1α* mediated protection against senescence in neonatal BMDMs.

We would like to highlight that our study is not without limitations. For example, although we have uncovered distinct differences in the secretomes and phenotypes of neonatal and adult BMDMs under different pathophysiologic stimuli, our study is descriptive and not causal in nature. While our findings are hypothesis generating, future mechanistic studies are required to answer why neonatal *HIF-1α* expressing BMDMs respond differently than adult *HIF-1α* expressing BMDMs to hypoxic stress and LPS stimulation. Secondly, we did not examine the proteomes of *HIF-1α* deficient adult BMDMs, and thus cannot comment on differences in *HIF-1α* dependent protein secretion to the aforementioned stimuli in adult BMDMs. Third, as late fetal and early neonatal myeloid seeding of the bone marrow is supplied primarily by the fetal liver, it is possible that the specific phenotype we observed in wildtype neonatal BMDMs is due to the embryonic, tissue resident-like nature of neonatal BMDMs and lost in adulthood due to replenishment of the bone marrow myeloid compartment by a non-embryonic-like progenitor [7,8]. Thus, in future studies, lineage tracing and fate mapping should be interrogated to answer whether the neonatal BMDM phenotype we observed is lost due to replacement of the neonatal bone marrow by a non-embryonic cell type versus epigenetic changes in HIF-1α targets. Finally, our study utilized an *in vitro*, controlled system in murine species to examine differences in the proteomes of neonatal and adult BMDMs under hypoxic stress and/or LPS stimulation. Thus, future investigations are required to confirm our proteomic findings in both murine and human species *in vivo* using pathophysiologic models, such as sepsis, to interrogate the clinical relevance of our findings.

In summary, we have newly identified distinct, age-specific differences in the secretomes and phenotypes of neonatal and adult BMDMs in response to common physiologic stressors such as hypoxia and LPS. Neonatal BMDMs assume an anti-inflammatory, regenerative phenotype protective against apoptosis and oxidative stress dependent on *HIF-1α*, while adult BMDMs adopt a pro-inflammatory, glycolytic phenotype focused on pathogen killing in response to hypoxic stress or LPS stimulation. How and when neonatal BMDMs transition to assume an adult-like response to these stimuli requires further investigation. Mechanistic interrogation of neonatal *HIF-1α* responses to these stimuli may also provide new targets to promote regenerative and anti-inflammatory programming of non-neonatal BMDMs to provide clinical benefit in sepsis and other disease states.

## Acknowledgements

This work was supported by funding through Northwestern University Clinical & Translational Sciences (NUCATS) Institute and the National Institutes of Health’s National Center for Advancing Translational Sciences KL2 grant award to A.B. (KL2TR001424) and funding from Lurie Children’s Hospital of Chicago to Thorp. Proteomics services were performed by the Northwestern Proteomics Core Facility, generously supported by NCI CCSG P30 CA060553 awarded to the Robert H Lurie Comprehensive Cancer Center, instrumentation award (S10OD025194) from NIH Office of Director, and the National Resource for Translational and Developmental Proteomics supported by P41 GM108569.

## Author Contributions

Conceptualization, A.B. and E.B.T.; Methodology, A.B. and M.F.; Investigation: A.B., M.F., C.L., and K.G; Software, A.B. and M.F.; Resources, A.B. and E.B.T.; Writing Original Draft, A.B. and E.B.T.; Review and Editing, all authors; Funding Acquisition, A.B. and E.B.T., Supervision, E.B.T.

## Declaration of Generative Artificial Intelligence (AI) in Scientific Writing

No generative AI or AI-assisted technologies were utilized in the writing process.

## Data Sharing Statement

Mass spectrometry raw data will be made immediately available upon request.

## Key Resources

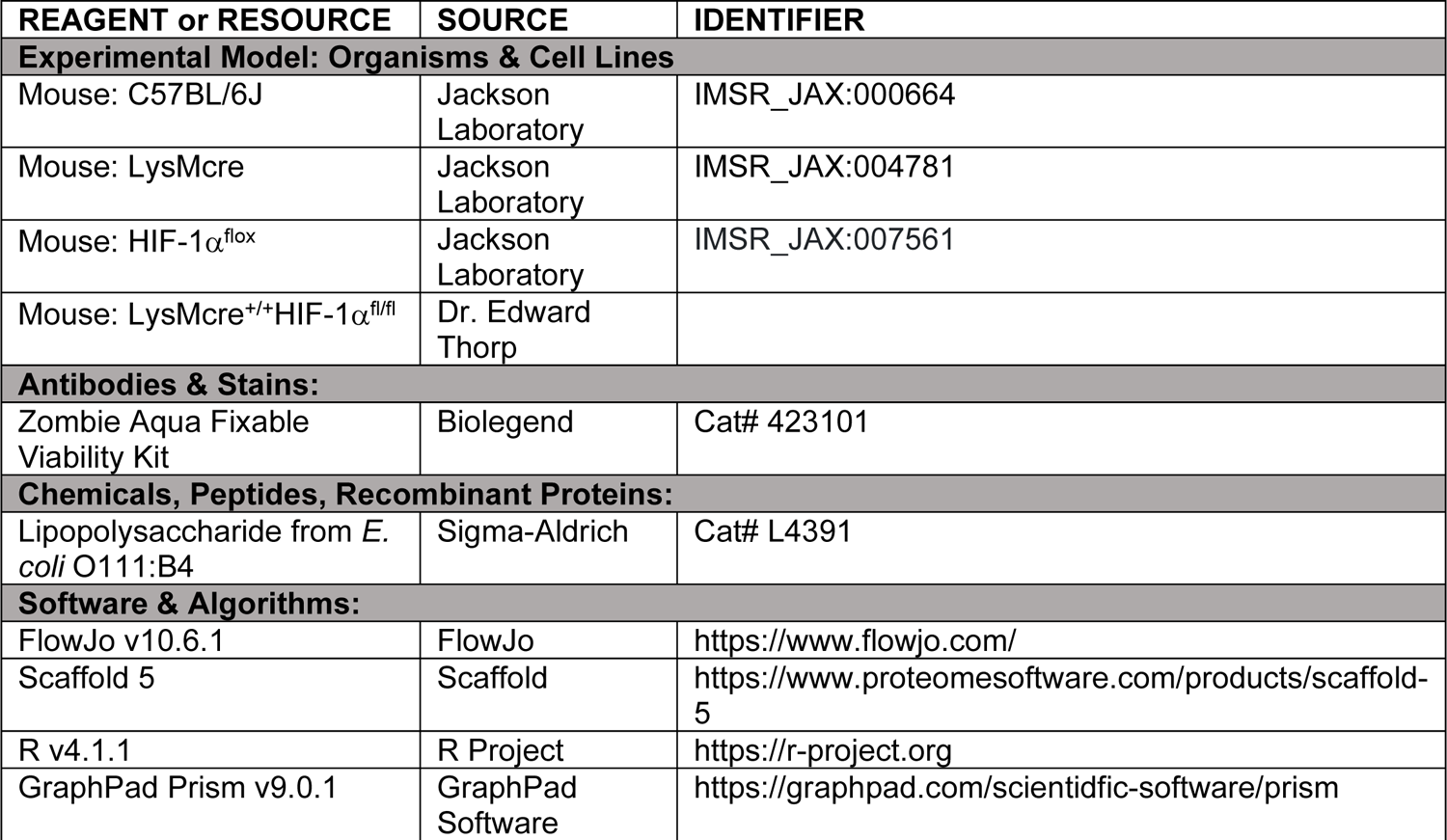

## Supplemental Figures

**Supplemental Figure 1:**
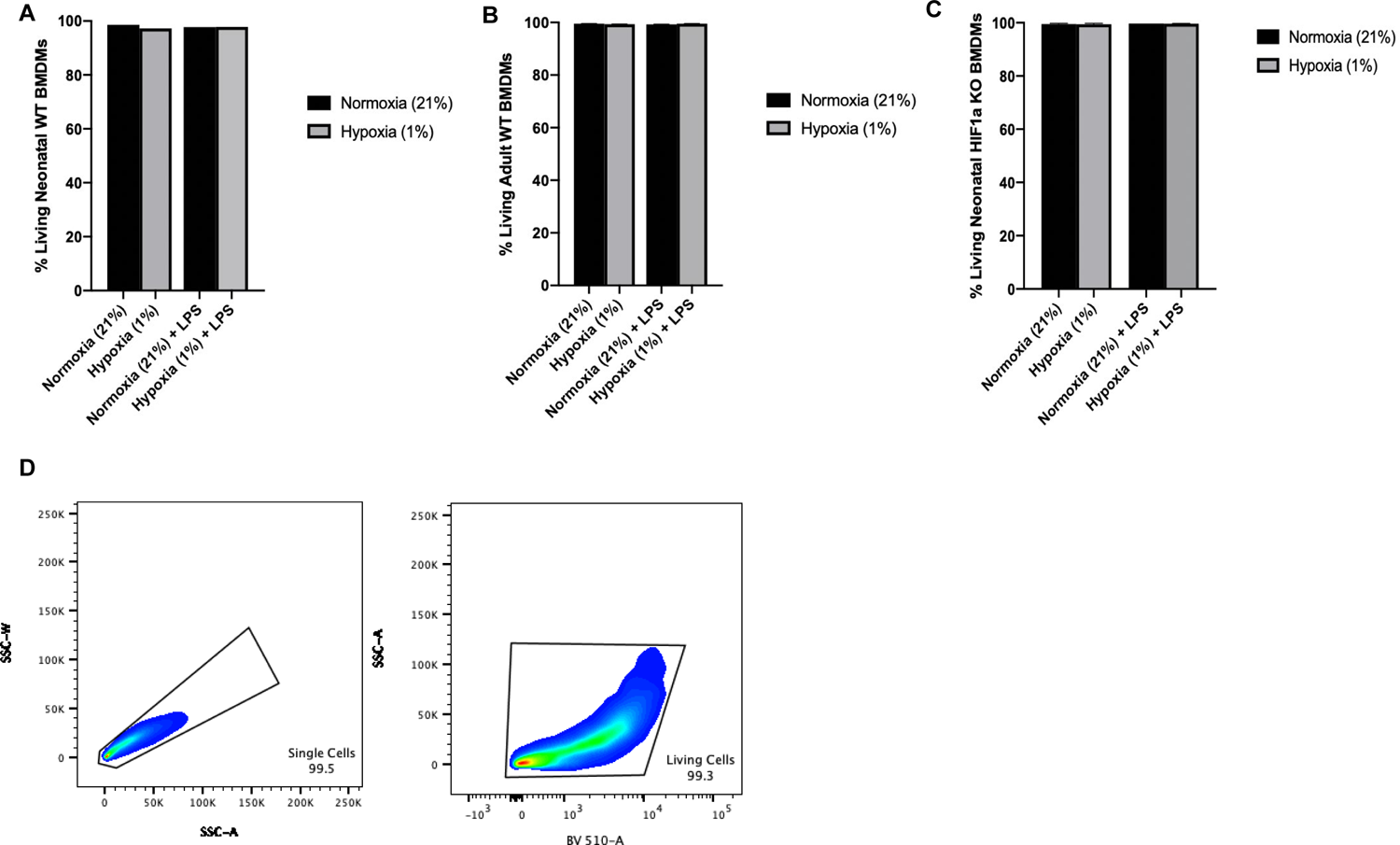
Neonatal and adult macrophage survival was not significantly impacted by hypoxia or LPS stimulation. Quantification of macrophage survival (live/dead staining) by flow cytometry in the setting of normoxia, hypoxia, normoxia + LPS, and hypoxia + LPS for (A) Neonatal wildtype BMDMs, (B) Adult wildtype BMDMs, and (C) Neonatal *LysMcre^+/+^HIF-1α^fl/fl^* (Neo HIF-1α KO) BMDMs. (D) Gating strategy for determination of living BMDMs.

**Supplemental Figure 2:**
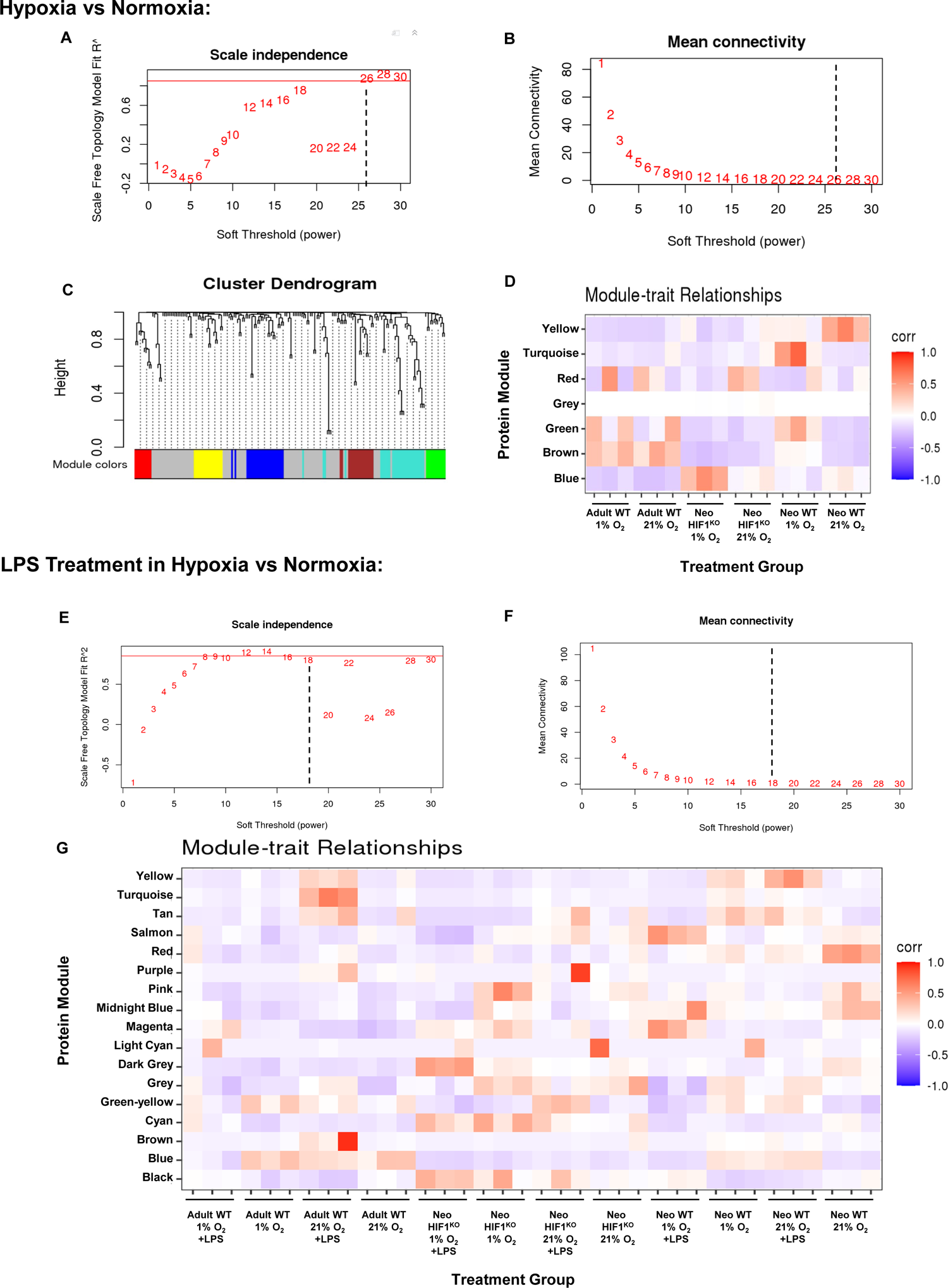
Protein cluster identification by weighted correlation network analyses (WCNA). (A) Graph showing the relationship between soft power and scale free topology model R^2^ for WCNA of normoxia versus hypoxia where the dashed line represents the soft power value chosen. (B) Graph showing the relationship between soft power and mean connectivity for WCNA of normoxia versus hypoxia where the dashed line represents the soft power value chosen. Soft power was chosen at the minimum value which resulted in a scale free topology model R^2^ > 0.85 with mean connectivity minimized. (C) Cluster dendrogram showing hierarchical clustering of proteins within the modules from WCNA of normoxia versus hypoxia. (D) Correlation heatmap showing how each sample from normoxia and hypoxia correlate to each protein module from WCNA. (E) Graph showing the relationship between soft power and scale free topology model R^2^ for WCNA analysis of normoxia and hypoxia with LPS treatment where the dashed line represents the soft power value chosen. (F) Graph showing the relationship between soft power and mean connectivity for WCNA analysis of normoxia and hypoxia with LPS treatment where the dashed line represented the soft power value chosen. Soft power was chosen at the minimum value which resulted in a scale free topology model R^2^ > 0.85 with mean connectivity minimized. (G) Correlation heatmap showing how each samples from normoxia and hypoxia with LPS treatment correlate to each protein module from WCNA.

**Supplemental Figure 3:**
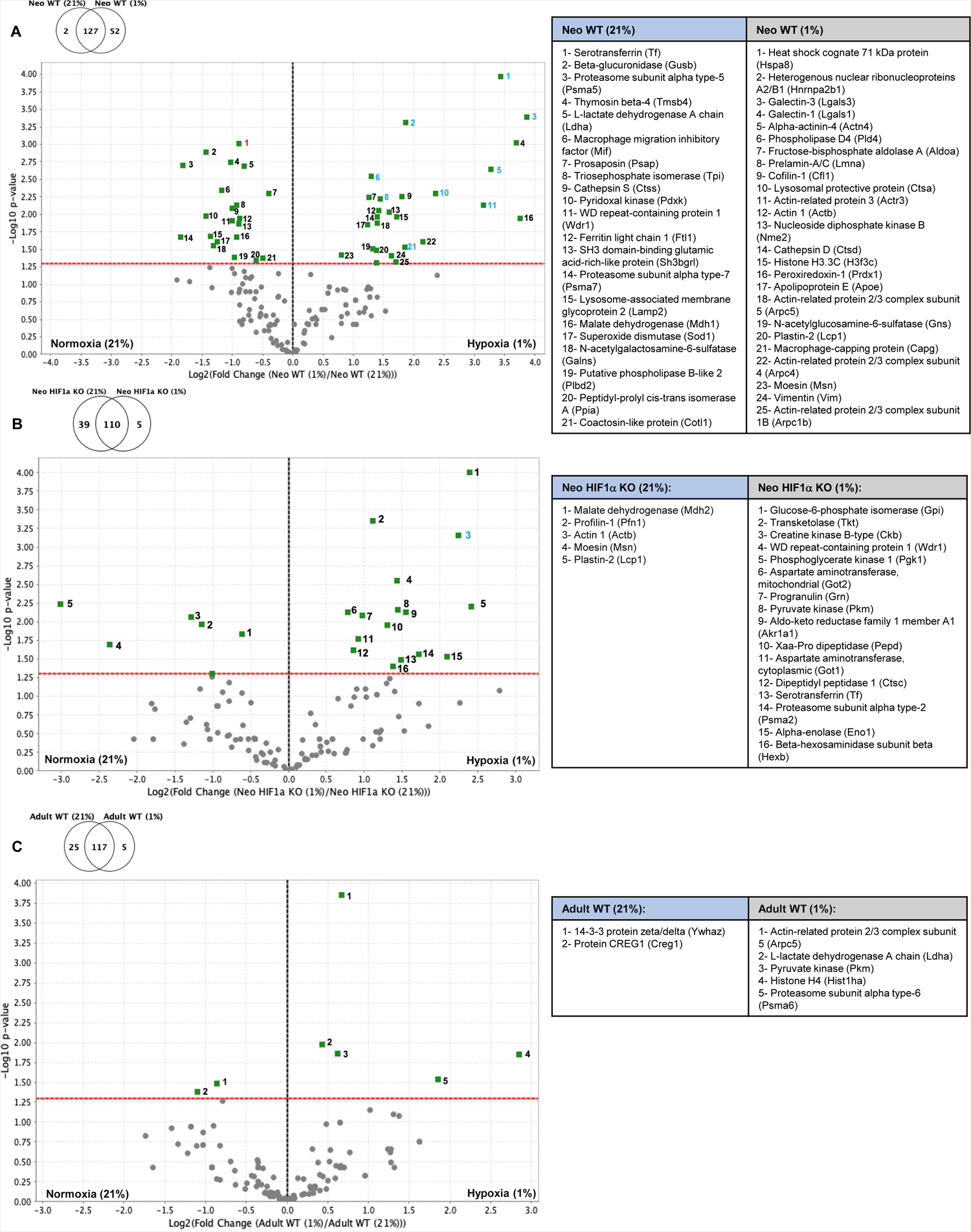
Hypoxia stimulates neonatal wildtype (Neo WT), neonatal *LysMcre^+/+^HIF-1α^fl/fl^* (Neo HIF-1α KO), and adult wildtype (Adult WT) macrophages to secrete significantly different proteins compared to normoxic conditions. Volcano plots depicting significant differences in protein secretion for (A) Neo WT BMDMs exposed to FiO2 0.21 vs FiO2 0.01 (B) Neo HIF-1α KO BMDMS exposed to FiO2 0.21 vs FiO2 0.01 (C) Adult WT BMDMs exposed to FiO2 0.21 vs FiO2 0.01. For all volcano plots the red dashed line represents the significance threshold and the black dashed line represents zero-fold change between groups utilizing t-tests with a significance level of < 0.05. Protein colored numbering: black= proteins secreted under both normoxic and hypoxic conditions, red= proteins only secreted under normoxic conditions, blue= proteins only secreted under hypoxic conditions.

**Supplemental Figure 4:**
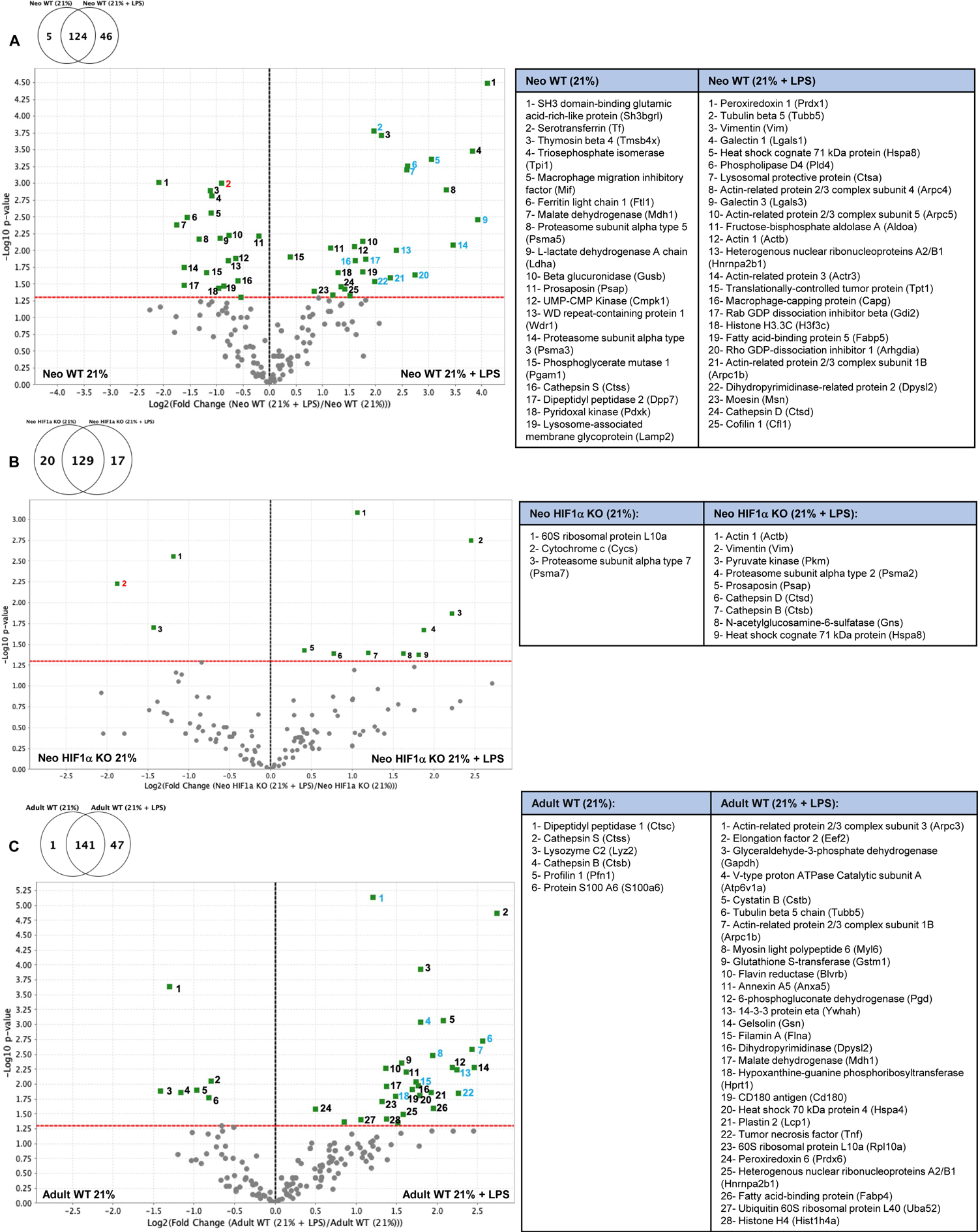
LPS stimulates neonatal wildtype (Neo WT), neonatal *LysMcre^+/+^HIF-1α^fl/fl^* (Neo HIF-1α KO), and adult wildtype (Adult WT) macrophages to secrete significantly different proteins compared to normoxia alone. Volcano plots depicting significant differences in protein secretion for (A) Neo WT BMDMs exposed to FiO2 0.21 vs FiO2 0.21+LPS (B) Neo HIF-1α KO BMDMS exposed to FiO2 0.21 vs FiO2 0.21+LPS (C) Adult WT BMDMs exposed to FiO2 0.21 vs FiO2 0.21+LPS. For all volcano plots the red dashed line represents the significance threshold and the black dashed line represents zero-fold change between groups utilizing t-tests with a significance level of < 0.05. Protein colored numbering: black= proteins secreted under both normoxic and LPS conditions, red= proteins only secreted under normoxic conditions, blue= proteins only secreted with LPS stimulation.

**Supplemental Figure 5:**
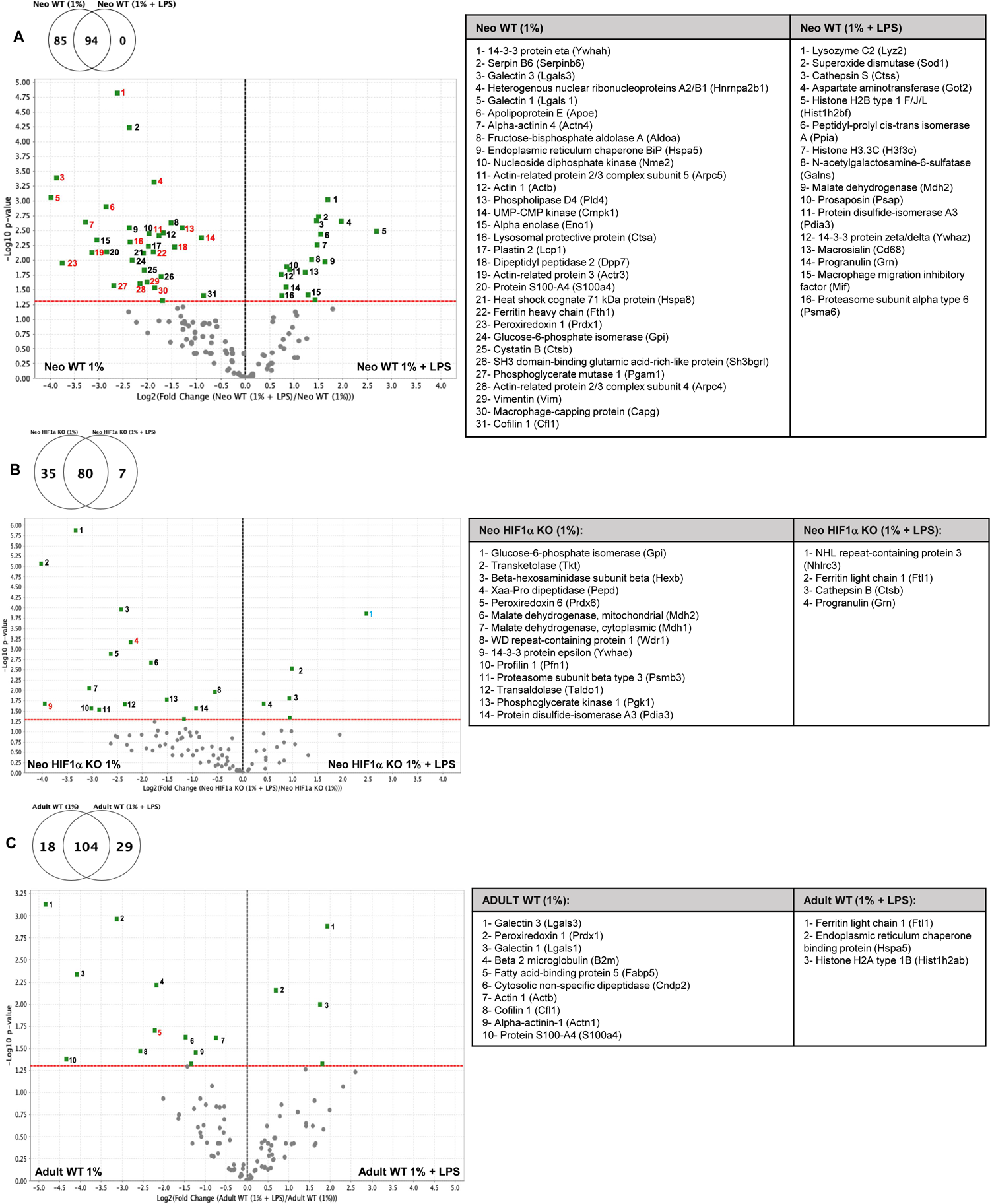
Hypoxia plus LPS stimulates neonatal wildtype (Neo WT), neonatal *LysMcre^+/+^HIF-1α^fl/fl^* (Neo HIF-1α KO), and adult wildtype (Adult WT) macrophages to secrete significantly different proteins compared to hypoxic stimulation alone. Volcano plots depicting significant differences in protein secretion for (A) Neo WT BMDMs exposed to FiO2 0.01 vs FiO2 0.01+LPS (B) Neo HIF-1α KO BMDMS exposed to FiO2 0.01 vs FiO2 0.01+LPS (C) Adult WT BMDMs exposed to FiO2 0.01 vs FiO2 0.01+LPS. For all volcano plots the red dashed line represents the significance threshold and the black dashed line represents zero-fold change between groups utilizing t-tests with a significance level of < 0.05. Protein colored numbering: black= proteins secreted with hypoxic stimulation alone and hypoxia + LPS, red= proteins only secreted under hypoxic conditions, blue= proteins only secreted with hypoxia + LPS stimulation.

**Supplemental Figure 6:**
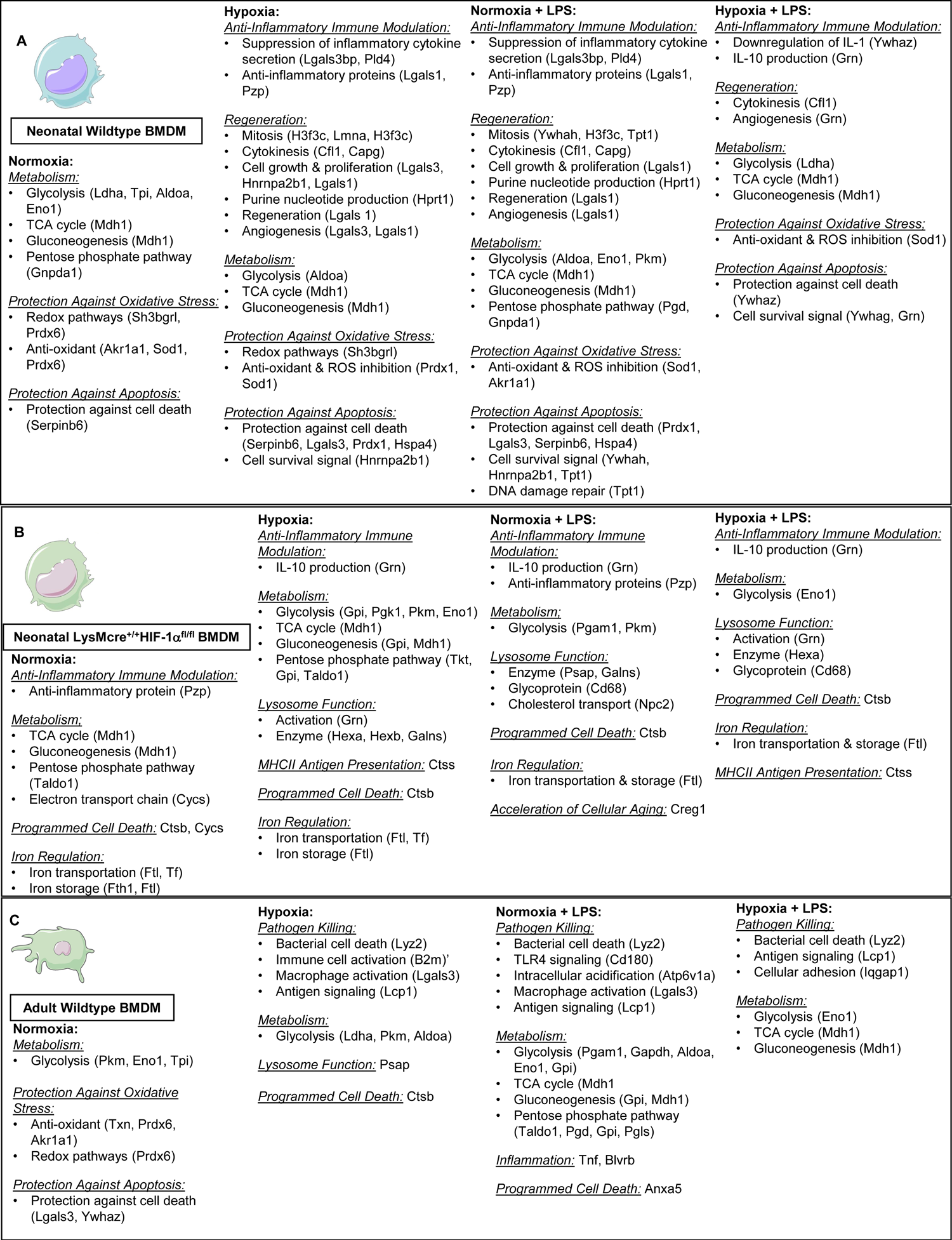
Summary of key differences in secreted proteomes from (A) neonatal wildtype, (B) neonatal *LysMcre^+/+^HIF-1a^fl/fl^*, and (C) adult WT BMDMs in response to normoxia, hypoxia, normoxia + LPS, and hypoxia + LPS.

---

**Supplemental Table 1:** Previously described biological functions of proteins secreted from neonatal wildtype, neonatal *LysMcre^+/+^HIF-1α^fl/fl^*, and adult WT BMDMs. Displayed proteins with previously described biological functions and references represent the significantly secreted proteins from neonatal and adult BMDMs based on volcano plot data utilizing t-tests with a significance level of < 0.05.

| Protein | Previously Described Biological Function | References |
| --- | --- | --- |
| 6-phosphogluconate dehydrogenase (Pgd) | Pentose phosphate pathway | 34 |
| 6-phosphogluconolactonase (Pgls) | Pentose phosphate pathway | 34 |
| 14-3-3 protein eta (Ywhah) | Signaling protein involved in cell cycle control, mitogenic signal transduction, cell survival | 35, 36 |
| 14-3-3 protein epsilon (Ywhae) | Signaling protein involved in cell cycle control, mitogenic signal transduction, cell survival | 36, 37 |
| 14-3-3 protein gamma (Ywhag) | Cell survival signal | 36, 38 |
| 14-3-3 protein zeta/delta (Ywhaz) | Signaling protein involved in cell survival, cell cycle progression, inhibition of apoptosis, downregulation of IL-1; activates autophagy under hypoxic stress; HIF-1a stabilization | 36, 38, 39 |
| 40S ribosomal protein S25 (Rps25) | Ribosome component; protein synthesis | 34 |
| 60S ribosomal protein L10a (Rpl10a) | Ribosome component; protein synthesis | 34 |
| Actin-1 (Actb) | Cytoskeleton (microfilament bundles, adherens-type junctions); cell motility, cell division/cytokinesis, vesicle and organelle movement, cell signaling | 34, 40, 41 |
| Actin-related protein 2/3 complex subunit 1B (Arpc1b) | Actin polymerization | 34 |
| Actin-related protein 2/3 complex subunit 3 (Arpc3) | Actin polymerization | 34 |
| Actin-related protein 2/3 complex subunit 4 (Arpc4) | Actin polymerization | 34 |
| Actin-related protein 2/3 complex subunit 5 (Arpc5) | Actin polymerization | 34 |
| Actin-related protein 3 (Actr3) | Actin polymerization | 34 |
| Aldo-keto reductase family 1 member A1 (Akr1a1) | NADPH-dependent reduction of aldehydes and ketones; resistance to oxidative stress; detoxifying enzyme | 42, 43 |
| Alpha-actinin-1 (Actn1) | Cytoskeleton protein | 34 |
| Alpha-actinin-4 (Actn4) | Cytoskeleton protein | 34 |
| Alpha-enolase (Eno1) | Glycolysis; cell surface receptor on activated immune cells allowing for tissue invasion; heat shock protein | 34, 44, 45 |
| Aminopeptidase N (Anpep) | Peptide breakdown | 46 |
| Annexin A5 (Anxa5) | Apoptosis | 47 |
| Apolipoprotein E (ApoE) | Fat binding protein; cholesterol metabolism; lipid-mediated antigen presentation | 48, 49 |
| Aspartate aminotransferase (Got1 &2) | Amino acid metabolism | 50 |
| Beta-glucuronidase (Gusb) | Catalytic carbohydrate metabolism | 51 |
| Beta-hexosaminidase subunit alpha (Hexa) | Lysosome enzyme | 34 |
| Beta-hexosaminidase subunit beta (Hexb) | Lysosome enzyme | 34 |
| Beta-2-microglobulin (B2m) | Immune cell activation | 52 |
| Calreticulin (Calr) | Endoplasmic reticulum chaperone binding protein; protein folding quality control; calcium homeostasis | 34 |
| Cathepsin B (Ctsb) | Lysosomal cysteine protease involved in programmed cell death | 53 |
| Cathepsin D (Ctsd) | Protein degradation; activation/degradation of polypeptide hormones and growth factors | 54 |
| Cathepsin S (Ctss) | Lysosomal cysteine protease involved in degradation of antigenic proteins for MHCII presentation; extracellular matrix cleavage | 34, 55 |
| Cathepsin Z (Ctsz) | Lysosomal cysteine protease | 34 |
| CD180 antigen (Cd180) | Toll-like receptor 4 (TLR4) signaling | 56 |
| Cofilin-1 (Cfl1) | Actin filament turnover; cell motility; cytokinesis | 57 |
| Coactosin-like protein (Cotl1) | Actin cytoskeleton; stabilizes 5-lipoxygenase (1st committed enzyme in arachidonic acid metabolism) | 34, 58 |
| Copper transport protein ATOX1 (Atox1) | Copper transport/homeostasis; anti-oxidant | 34 |
| Creatine kinase B-type (Ckb) | Catalyzes transfer of phosphate between ATP; inhibits ferroptosis | 34, 59 |
| Cystatin-B (Cstb) | Inhibits cathepsins; prevents lysosomal protease leakage | 34 |
| Cytosolic non-specific dipeptidase (Cndp2) | Hydrolyzes dipeptides | 60 |
| Cytochrome C (CyCs) | Mitochondrial electron transport chain; apoptosis; ferrous/ferric conversion | 61, 62 |
| Dihydropyrimidinase-related protein 2 (Dpysl2) | Microtubule assembly | 34 |
| Dipeptidyl peptidase 1 (Dpp1) | Serine exopeptidase that cleaves dipeptides from the N-terminus of polypeptides; cleaves growth factors, chemokines, neuropeptides, and vasoactive peptides; increases serum glucose | 34 |
| Dipeptidyl peptidase 2 (Dpp7) | Serine exopeptidase that cleaves dipeptides from the N-terminus of polypeptides; cleaves growth factors, chemokines, neuropeptides, and vasoactive peptides; increases serum glucose | 34 |
| Elongation factor 2 (Eef2) | Protein synthesis; facilitates GTP-dependent translocation of the ribosome | 34 |
| Endoplasmic reticulum chaperone binding protein (Hspa5) | Chaperones newly synthesized proteins; maintains permeability of the endoplasmic reticulum; retrograde translocation of misfolded proteins for degradation by the proteasome | 34, 63 |
| Fatty acid-binding protein (Fabp4) | Fatty acid transport | 34 |
| Fatty acid-binding protein 5 (Fabp5) | Fatty acid transport | 34 |
| Ferritin light chain 1 (Ftl) | Iron transportation & storage; iron detoxification | 34, 64 |
| Ferritin heavy chain (Fth1) | Intracellular iron storage | 34 |
| Filamin-A (Flna) | Cytoskeleton formation; integrin binding | 65 |
| Flavin reductase (Blvrb) | Catalyzes NADPH dependent reactions (methemoglobin); H2O2 production | 34 |
| Fructose-bisphosphate aldolase A (Aldoa) | Glycolysis | 34 |
| Galectin-1 (Lgals1) | Autocrine growth factor that regulates cell proliferation (proliferative and antiproliferative- [dependent]); anti and pro-inflammatory; angiogenesis; T-cell regulation; regeneration | 66, 67 |
| Galectin-3 | Cell to cell communication; cell growth/differentiation; macrophage activation; angiogenesis; protection from apoptosis | 68, 69, 70 |
| Galectin-3-binding protein (Lgals3bp) | Suppresses inflammatory cytokine secretion | 71 |
| Gelsolin (Gsn) | Actin filament assembly and disassembly | 34 |
| Glucosamine-6-phosphate isomerase 1 (Gnpda1) | Pentose phosphate pathway | 34 |
| Glucose-6-phosphate isomerase (Gpi) | Glycolysis; gluconeogenesis; pentose phosphate pathway | 72, 73 |
| Glutathione reductase (Gsr) | Catalyzes reduction of glutathione disulfide | 74 |
| Glutathione S-transferase (Gstm1) | Glutathione conjugation for cell detoxification; inhibits apoptosis | 75 |
| Glyceraldehyde-3-phosphate dehydrogenase (Gapdh) | Glycolysis | 34 |
| Heat Shock 70 kDa protein 4 (Hspa4) | Inhibition of autophagy/apoptosis; cell growth/differentiation; correct protein folding | 76, 77 |

Supplemental Table 1 Continued
| Protein | Previously Described Biological Function | References |
| --- | --- | --- |
| Heterogenous nuclear ribonucleoproteins A2/B1 (Hnrnpa2b1) | RNA metabolism; cell survival; cell proliferation | 78, 79 |
| Histone H2A type 1B (Hist1h2ab) | DNA packaging into chromatin; epigenetic DNA modification | 80, 81 |
| Histone H2B type 1 F/J/L (Hist1h2bf) | DNA organization; epigenetic regulation of transcription | 80, 81 |
| Histone H3.3C (H3f3c) | Gene transcription; genomic stabilization; mitosis | 80, 81, 82 |
| Histone H4 (Hist1h4a) | Chromosome structure & function | 80, 81 |
| Hypoxanthine-guanine phosphoribosyltransferase (Hprt1) | Purine nucleotide production | 34 |
| L-lactate dehydrogenase A chain (Ldha) | Glycolysis | 34 |
| Lysosome-associated membrane glycoprotein 1 (Lamp1) | Lysosome membrane protein | 83 |
| Lysosome-associated membrane glycoprotein 2 (Lamp2) | Lysosome membrane protein | 83 |
| Lysosomal protective protein (Ctsa) | Stabilizes lysosomal enzymes | 34 |
| Lysozyme C-2 (lyz2) | Bacterial cell death; protein catabolism | 84 |
| Macrophage-capping protein (Capg) | Regulation of actin polymerization necessary for proper cytokinesis | 85 |
| Macrophage migration inhibitory factor (Mif) | Pro-inflammatory protein; upregulates expression of TLR4; inhibits p53-dependent apoptosis of macrophages; counter regulates the immunosuppressive effects of glucocorticoids on immune cells | 34, 86 |
| Macrosialin (Cd68) | Lysosome/endosome glycoprotein | 34 |
| Malate dehydrogenase (Mdh1) | TCA cycle; gluconeogenesis | 34, 87 |
| Moesin (Msn) | Cross links plasma membranes and actin cytoskeletons | 34 |
| Myosin light polypeptide 6 (MyI6) | Motor protein | 34 |
| N-acetylgalactosamine-6-sulfatase (Galns) | Lysosome enzyme | 34 |
| NHL repeat-containing protein 3 (Nhlrc3) | Protein ubiquination | 34 |
| NPC intracellular cholesterol transporter 2 (Npc2) | Lysosome/endosome cholesterol transport | 34 |
| Nucleoside diphosphate kinase B (Nme2) | Catalyzes exchange of terminal phosphate between nucleoside diphosphates and triphosphates | 88 |
| Peptidyl-prolyl cis-trans isomerase A (Ppia) | Catalyzes the cis/trans isomerization of the peptidyl-prolyl peptide bond in oligopeptides and proteins, a rate limiting step in the process of protein folding to generate functional proteins | 34 |
| Peroxiredoxin 6 (Prdx6) | Anti-oxidant; redox regulation | 34, 89 |
| Peroxiredoxin 1 (Prdx1) | Anti-oxidant/ROS inhibitor; cell growth/differentiation; apoptosis inhibitor | 34, 90 |
| Phosphoglycerate kinase 1 (Pgk1) | Glycolysis | 34 |
| Phosphoglycerate mutase 1 (Pgam1) | Glycolysis | 34 |
| Phospholipase D4 (Pld4) | Hematopoietic progenitor cell differentiation; cytokine regulation | 34 |
| Plastin-2 (Lcp1) | Cross-links actin filaments into tight bundles to facilitate antigen receptor signaling, cell adhesion, and motility | 91 |
| Pregnancy zone protein (Pzp) | Protease inhibitor; anti-inflammatory immune modulation; fibrinolysis | 92, 93, 94 |
| Prelamin-A/C (Lmna) | Mitosis; nuclear organization and regulation of nuclear processes (DNA replication, transcription, and chromatin organization) | 34, 95 |
| Profilin-1 (Pfn1) | Actin polymerization | 34 |
| Progranulin (Grn) | Cell survival; angiogenesis; IL-10 production; lysosome activation/function | 96, 97, 98, 99 |
| Prosaposin (Psap) | Lysosome function | 34 |
| Proteasome subunit alpha type-2 (Psmα2) | Proteolysis | 34 |
| Proteasome subunit alpha type-3 (Psmα3) | Proteolysis | 34 |
| Proteasome subunit alpha type-5 (Psmα5) | Proteolysis | 34 |
| Proteasome subunit alpha type-6 (Psmα6) | Proteolysis | 34 |
| Proteasome subunit alpha type-7 (Psmα7) | Proteolysis | 34 |
| Proteasome subunit beta type-3 (Psmβ3) | Proteolysis | 34 |
| Proteasome subunit beta type-8 (Psmβ8) | Proteolysis | 34 |
| Protein disulfide-isomerase A3 (Pdia3) | Mediates proper folding of glycoproteins | 34 |
| Protein CREG1 (Creg1) | Accelerates cellular senescence; autophagy; inhibitors proliferation; promotes differentiation | 100, 101, 102, 103 |
| Protein S100 (S100a4) | Cell motility; cell invasion; cell survival; cell cycle progression and differentiation; angiogenesis | 34, 104 |
| Putative phospholipase B-like 2 (Plbd2) | Lipid catabolism | 34 |
| Pyridoxal kinase (Pdxk) | Enzyme required to convert B6 to its active form | 34 |
| Pyruvate kinase (Pkm) | Glycolysis | 34 |
| Rab GDP dissociation inhibitor beta (Gdi2) | Inhibits GDP to GTP exchange; regulates vesicular trafficking of molecules between cells; NADPH oxidase inactivation | 34 |
| Ras GTPase-activating-like protein IQGAP1 (Iqgap1) | Scaffold protein involved in organization of the cytoskeleton and cellular adhesion | 105 |
| Reticulon-4 (Rtn4) | Inhibition of neuroendocrine signal secretion; vascular remodeling | 106, 107 |
| Rho GDP-dissociation inhibitor 1 (Arhgdia) | Inhibits GDP to GTP exchange; regulates vesicular trafficking of molecules between cells; NADPH oxidase inactivation | 108 |
| Serpin B6 (Serpinb6) | Cell death inhibitor | 109 |
| Serotransferrin (Tf) | Iron transportation & metabolism | 110 |
| SH3 domain-binding glutamic acid-rich-like protein (Sh3bgrl) | Protection against oxidative stress (redox pathways) | 111 |
| Superoxide dismutase (Sod1) | ROS inhibitor & anti-oxidant | 34 |
| Thioredoxin (Txn) | Anti-oxidant | 34 |
| Thymosin beta-4 (Tmsb4x) | Angiogenesis; wound healing; cell proliferation, migration, and differentiation | 34, 112, 113 |
| Transaldolase (Taldo1) | Pentose phosphate pathway | 34 |
| Transketolase (Tkt) | Pentose phosphate pathway: NADPH production to combat oxidative stress | 34 |
| Translationally-controlled tumor protein (Tpt1) | Cell survival; DNA damage repair; mitosis | 34, 114 |
| Transmembrane glycoprotein (Gpnmb) | Cell-cell recognition | 115 |
| Triosephosphate isomerase (Tpi) | Glycolysis | 34 |
| Tubulin beta-5 chain (Tubb5) | Structural component of the cytoskeleton involved in the regulation of synapse organization; microtubule formation | 34 |

Supplemental Table 1 Continued
| Protein | Previously Described Biological Function | References |
| --- | --- | --- |
| Tumor necrosis factor (Tnf) | Adipokine (insulin resistance); cytokine (immune cell regulation, pro-inflammatory, apoptosis) | 116, 117 |
| Ubiquitin 60S ribosomal protein L40 (Uba52) | Protein degradation | 34 |
| UMP-CMP kinase (Cmpk1) | Nucleic acid biosynthesis | 34 |
| Vimentin (Vim) | Microtubule and cytoskeleton stabilization | 34 |
| V-type proton ATPase catalytic subunit A (Atp6v1a) | Acidification of intracellular organelles | 118 |
| WD repeat-containing protein 1 (Wdr1) | Disassembly of actin filaments | 34 |
| Xaa-Pro Dipeptidase (Pepd) | Protein catabolism | 34 |

